# Simulated Solute Tempering 2: An Efficient and Practical Approach to Protein Conformational Sampling and Binding Events

**DOI:** 10.1101/2024.10.03.613476

**Authors:** Dirk Stratmann, Gautier Moroy, Pierre Tuffery, Samuel Murail

## Abstract

Molecular dynamics (MD) simulations are powerful tools for studying the movement and interactions of molecules, but they can be computationally expensive, especially for large biomolecules like proteins. This is problematic because accurately simulating the motions of these molecules is key to understanding their function. Enhanced sampling methods, such as Simulated Tempering (ST), temperature Replica Exchange Molecular Dynamics (REMD) and Replica Exchange with Solute Tempering (REST and REST2), have been developed to overcome this challenge by improving the efficiency of MD simulations. This article presents a new enhanced sampling method called Simulated Solute Tempering 2 (SST2) that builds upon the strengths of ST and REST2. SST2 selectively scales the interactions inside a biomolecule and with its surrounding environment, effectively accelerating the exploration of its different structural states and their stability at various temperatures. SST2 was tested on three different systems (chignolin CLN025, Trp-Cage, and a protein-peptide complex, p97/PNGase) and found to achieve comparable or superior sampling efficiency to ST, SST1 and REST2 while requiring fewer temperature rungs. Notably, SST2 is particularly well-suited for investigating large biomolecular systems, making it a valuable tool for studying a wide range of biomolecular processes, from protein folding to ligand binding.

**Significance Statement:** Accurately simulating the motions of biological molecules like proteins is key to understanding their function. However, this is computationally expensive, especially for large systems or slow processes. To overcome this, we developed Simulated Solute Tempering 2 (SST2), a novel algorithm that improves the efficiency of molecular dynamics (MD) simulations. SST2 selectively scales the interactions inside a biomolecule and with its surrounding environment, accelerating the exploration of its different structures and their stability at various temperatures. This allows SST2 to achieve remarkable performances with a clear improved ease of use compare to similar methods, particularly for large systems like protein-ligand complexes. This makes SST2 a valuable tool for studying a wide range of biomolecular processes, from protein folding to ligand binding.

**TOC Graphic:** 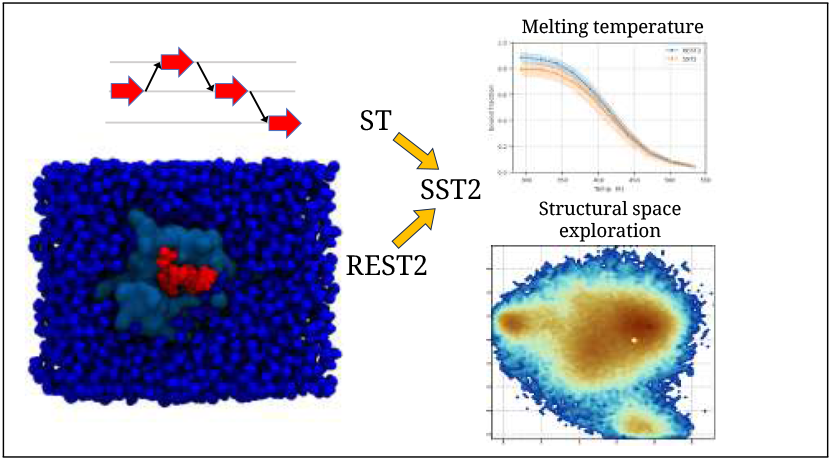

## Introduction

Recent breakthroughs in deep learning, exemplified by AlphaFold 2, ^1^ have led to unprecedented success in protein structure prediction. This progress represents a crucial step towards a deeper understanding of protein function. Derivatives of AlphaFold 2, such as Colabfold^2^ and AlphaFold-Multimer, ^3^ have further extended the scope of protein structure prediction, allowing the modeling of protein complexes and multimers. In the context of protein complexes, the specific case of proteinpeptide complexes has demonstrated that these methods can accurately predict the structure of protein-peptide complexes for 60 to up to 90 % of complexes. ^4,5^ However, predicting a static protein structure is only one part of the story, and a nuanced understanding of protein function requires a thorough exploration of protein dynamics behavior and the multiple conformations that proteins can adopt. Molecular dynamics (MD) simulations, based on Newton’s equations of motion, provide a means of predicting the temporal evolution of biomolecular systems.^6–8^ Despite their potential, MD simulations are often limited by their computational cost when it comes to capturing the timescales of critical conformational changes, particulary at the microsecond scale and beyond.

During MD simulations, proteins frequently become trapped in local minima of the free energy landscape, which hinders the exploration of conformational diversity. To overcome these limitations, several enhanced sampling techniques have been developed such as temperature Replica Exchange Molecular Dynamics (for simplicity REMD will be used as temperature REMD) ^9^ and Simulated Tempering (ST).^10,11^ REMD runs parallel simulations at different temperatures, allowing periodic exchanges between replicas based on a Metropolis criterion. In contrast, ST involves a single simulation with periodic updates of its temperature, guided by a Metropolis criterion that takes into account potential energy and weights.

While REMD and ST are effective in solving local minima problems, their efficiency depends on the size of the simulated system, as the optimal number of replicas or temperature ladders increases greatly with the degrees of freedom of the simulated system. For large systems, the number of replicas required to achieve efficient temperature sampling is too high to be practical.

In order to address these challenges, Liu *et al*. introduced Replica Exchange with Solute Tempering (REST).^12^ the REST method separates the system into solute and solvent, allowing the solute to ascend the temperature ladder while *maintaining* the solvent at ambient temperature (this is a physical approximation, the system is in fact warmed up, however in combination with the modified potential energy function, the solvent is considered to be maintained at ambient temperature). However, the REST method has been demonstrated to be less efficient than REMD for larger systems, such as Trp-Cage and *β* hairpin.^13^ The subsequent REST2 method^14^ is a pure Hamiltonian REMD. It ensures that all replicas run at the same temperature, but evolve with different Hamiltonians. Only the solute-solute and solute-solvent interactions are modified to mimick the effect of elevated temperature. The main difference between REST and REST2 is the scaling of the solute-solvent term. REST2 employs a weaker solute-solvent interaction at *“high temperature”*, which favoures the sampling of folded conformations at this conditions. Since then, REST2 has become a benchmark for conformational sampling of small proteins in different contexts.^15–17^ It has been implemented in popular MD simulation software like Gromacs and NAMD.^18–20^

In addition to its efficiency, REST2 has the advantage of simulating the solvent at ambient temperature, mitigating the challenges associated with the limited parametrisation of water molecules at elevated temperatures. This feature, demonstrated by Stirnemann and Sterpone, allows REST2 to more accurately reconstruct the thermal stability of proteins compared to REMD.^18^

It is worth noting that replica exchange, which depends on the potential energy of two simulations, has a lower probability of exchange compared to ST, resulting in a lower rate of traversal of energy space. ^21^ In addition, Rosta and Hummer^22^ have shown that running multiple ST simulations in parallel without communication is as efficient as running REMD simulations with communication for replica exchange. The need for communication between replicas implies the utilisation of distributed computations on CPU or GPU nodes that exhibit identical computational performance. As a result, both REMD and REST2 require specific and significant computing power. This can lead to significant performance limitations, particularly in scenarios where the CPU/GPU node park is heterogeneous. If computational performance varies between ladders, a simulation may need to wait for others to complete before attempting replica exchange and continuing the calculation.

The first version of Simulated Solute Tempering (SST1)^23^ method was developed as a combination of ST and REST. Like REST, SST1 allows a reduced number of replicas, and compared to ST, SST1 shows the highest round trip rate and the highest sampling speed on a alanine octapeptide. ^23^ However SST1 suffers from the same weaknesses of the first version of REST, and SST1 may be limited for the folding of larger systems with low exchange between folded and extended forms of proteins involving large conformational changes during folding.

In response to these challenges, we present a novel algorithm, Simulated Solvent Tempering 2 (SST2), designed to overcome the limitations of existing methods. By exploiting the strengths of ST and REST2, SST2 aims to achieve both efficient sampling and a straightforward implementation.

To assess the performance of SST2, extensive MD simulations were performed on three different systems, two *toy systems*, chignolin CLN025 and Trp-Cage and a protein-peptide complex, PNGase in complex with p97 peptide. Chignolin CLN025 and Trp-Cage are commonly used to benchmark protein folding methods.^18,24–28^ CLN025 is a *β* hairpin and TrpCage is an *α* helix. For the two *toy systems*, SST2 conformational space sampling capabilities have been systematicly compared to ST, SST1 and REST2. The ability of SST2 to reconstruct the thermal stability curve of simulated peptides has been investigated, and the effect of the reference temperature on the efficiency of SST2 has been studied. We also focused on the effect of *cis-trans* isomerisation of proline residues on sampling efficiency, and how to improve it.

Finally, to assess the algorithm sampling efficiency in ligand binding events and to study larger systems, whose size and complexity would prevent the use of REMD or ST, we applied SST2 to the peptide-protein complex, p97/PNGase and compare its performance against REST2.

## Results

### CLN025 and Trp-Cage simulations

As a first evaluation of the SST2 method, we performed explicit solvent simulations of two protein toy systems, chignolin CLN025 and Trp-Cage, starting from both folded (PDB IDs: 5AWL and 1L2Y) and unfolded conformations. For each system, we ran ST, SST1, REST2, and SST2 simulations (Table 1). ST simulations were performed with 20 temperature rungs between 280-500 *K*, while SST2 simulations used 10 *λ* rungs corresponding to effective solute-solute temperatures between 280-540 *K* at a reference solvent temperature of 300 K. Each trajectory was propagated for 10 *µs* (CLN025) or 40 *µs* (Trp-Cage), with four independent replicas for folded and unfolded states.

**Table 1:** List of simulation. PDB reference ID structure for of chignolin CLN025, Tryptophan Cage, PNGase-p97 and PNGase in free form is 5awl, ^29^ 1l2y,^30^ 2hpl and 2hpj, ^31^ respectively. All simulation were conducted with an exchange time of 2.0 *ps*.

| name | protein | starting form | length ($\mu s$) | friction ($ps^{-1}$) | Sampling Method | min. Temp. (K) | max. Temp. (K) | ref. Temp. (K) | #Rungs | Pro $\omega$ scaling | #Replicas |
| --- | --- | --- | --- | --- | --- | --- | --- | --- | --- | --- | --- |
| $F_{ST}$ | CLN025 | 5awl | 10.0 | 10.0 | ST | 280 | 500 | | 20 | | 4 |
| $U_{ST}$ | CLN025 | unfolded | 10.0 | 10.0 | ST | 280 | 500 | | 20 | | 4 |
| $F_{SST1}$ | CLN025 | 5awl | 10.0 | 10.0 | SST1 | 280 | 500 | 300 | 10 | | 4 |
| $U_{SST1}$ | CLN025 | unfolded | 10.0 | 10.0 | SST1 | 280 | 500 | 300 | 10 | | 4 |
| $F_{REST2}$ | CLN025 | 5awl | 1.0 | 1.0 | REST2 | 280 | 540 | 300 | 10 | yes | 4 |
| $U_{REST2}$ | CLN025 | unfolded | 1.0 | 1.0 | REST2 | 280 | 540 | 300 | 10 | yes | 4 |
| $F$ | CLN025 | 5awl | 10.0 | 1.0 | SST2 | 280 | 540 | 300 | 10 | yes | 4 |
| $U$ | CLN025 | unfolded | 10.0 | 1.0 | SST2 | 280 | 540 | 300 | 10 | yes | 4 |
| $F_{350K}$ | CLN025 | 5awl | 10.0 | 1.0 | SST2 | 280 | 540 | 350 | 10 | yes | 4 |
| $F_{6\text{ rungs}}$ | CLN025 | 5awl | 10.0 | 1.0 | SST2 | 280 | 540 | 300 | 6 | yes | 4 |
| $F_{big\ box}$ | CLN025 | 5awl | 10.0 | 1.0 | SST2 | 280 | 540 | 300 | 10 | yes | 2 |
| $F_{ST}$ | Trp-Cage | 1l2y | 40.0 | 10.0 | ST | 280 | 500 | | 20 | | 4 |
| $U_{ST}$ | Trp-Cage | unfolded | 40.0 | 10.0 | ST | 280 | 500 | | 20 | | 4 |
| $F_{SST1}$ | Trp-Cage | 1l2y | 20.0 | 10.0 | SST1 | 280 | 500 | 300 | 10 | | 4 |
| $U_{SST1}$ | Trp-Cage | unfolded | 20.0 | 10.0 | SST1 | 280 | 500 | 300 | 10 | | 4 |
| $F_{REST2}$ | Trp-Cage | 1l2y | 2.0 | 1.0 | REST2 | 280 | 540 | 300 | 10 | yes | 2 |
| $U_{REST2}$ | Trp-Cage | unfolded | 2.0 | 1.0 | REST2 | 280 | 540 | 300 | 10 | yes | 2 |
| $F$ | Trp-Cage | 1l2y | 40.0 | 1.0 | SST2 | 280 | 540 | 300 | 10 | yes | 2 (1,2) |
| $F$ | Trp-Cage | 1l2y | 40.0 | 1.0 | SST2 | 280 | 540 | 300 | 10 | no | 2 (3,4) |
| $U$ | Trp-Cage | unfolded | 40.0 | 1.0 | SST2 | 280 | 540 | 300 | 10 | yes | 2 (1,2) |
| $U$ | Trp-Cage | unfolded | 40.0 | 1.0 | SST2 | 280 | 540 | 300 | 10 | no | 2 (3,4) |
| $F_{350K}$ | Trp-Cage | 1l2y | 40.0 | 1.0 | SST2 | 280 | 540 | 350 | 10 | no | 4 |
| $F_{280-700K}$ | Trp-Cage | 1l2y | 20.0 | 1.0 | SST2 | 280 | 700 | 300 | 14 | yes | 4 |
| $F_{REST2}$ | PNGase-p97 | 2hpl | 1.0 | 1.0 | REST2 | 280 | 700 | 320 | 10 | | 4 |
| $U_{REST2}$ | PNGase-p97 | 2hpj + linear peptide | 1.0 | 1.0 | REST2 | 280 | 700 | 320 | 10 | | 4 |
| $F$ | PNGase-p97 | 2hpl | 20.0 | 1.0 | SST2 | 280 | 700 | 320 | 10 | | 4 |
| $U$ | PNGase-p97 | 2hpj + linear peptide | 20.0 | 1.0 | SST2 | 280 | 700 | 320 | 10 | | 4 |

For SST1, the simulation lengths were 10 *µs* (CLN025) and 20 *µs* (Trp-Cage), while REST2 simulations were run for 1 *µs ×* 10 replicas (CLN025) and 2 *µs ×* 10 replicas (Trp-Cage), using the same *λ* ladder as SST2. All simulations were performed with the Amber14SB force field, TIP3P water, and a 4 fs integration timestep with hydrogen mass repartitioning. Exchange attempts were carried out every 2 *ps*. More details on the simulation setup can be found in the Material and Methods section.

#### Transition acceptance ratio

With 10 *λ* rungs, SST2 simulations showed a high acceptance ratio energy 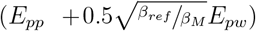 overlap for CLN025 (Fig. 1B) and Trp-Cage (Fig. S1B), resulting in an average probability of *λ* transition of about 84% and 66% for CLN025 and Trp-Cage, respectively (Fig. 1A and S1A). During ST simulations of CLN025 and Trp-Cage, we doubled the number of temperature rungs, resulting in a lower transition probability of about 47% and 31%, respectively (Fig. S3A and S4A).

**Figure 1.**
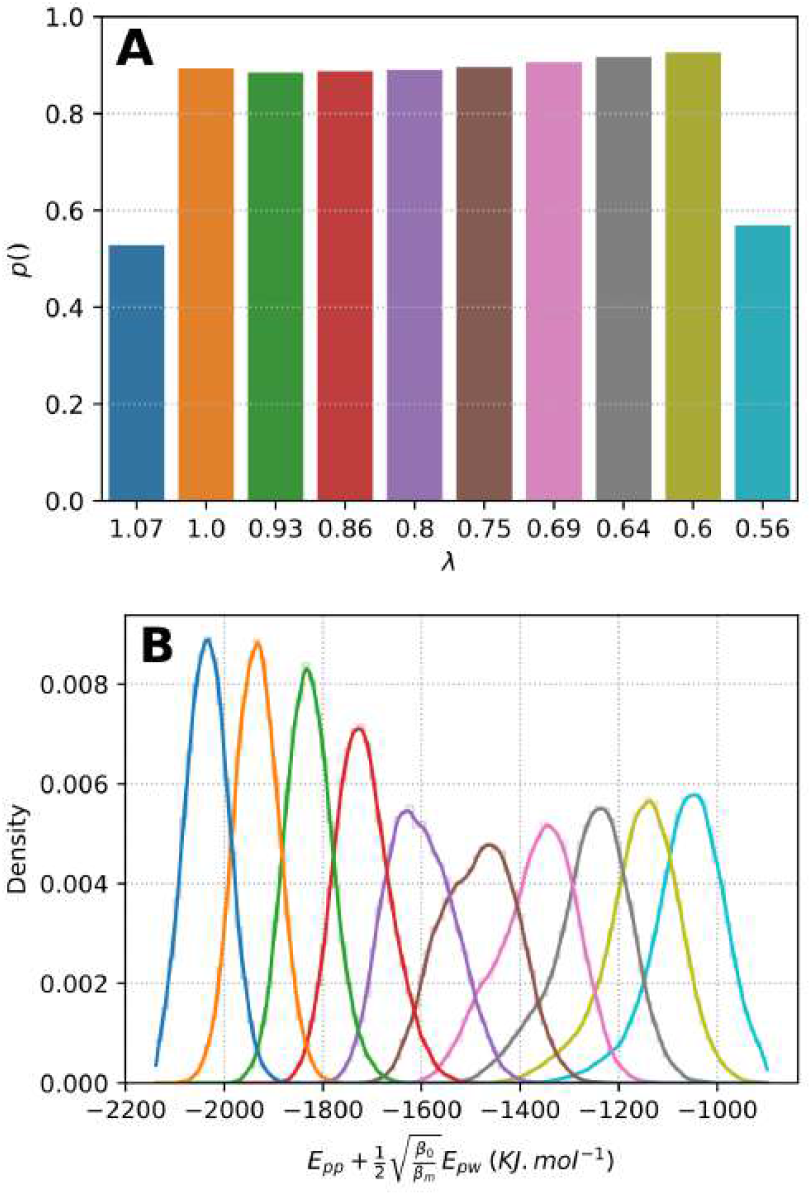
Probability of transition and distribution of the SST2 energy as function of *λ*_*i*_ for simulation CLN025 F(1). (A) Probability of transition at each *λ* rung. (B) Distribution of acceptance ratio energy 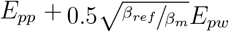 for each *λ*, the color code for *λ* value is identical to that of panel A. Maximum *λ* value 1.07 corresponds to a solute-solute temperature of 280 *K*, as minimum *λ* value 0.56, correspond to a temperature of 540 *K*.

For CLN025, we additionally investigated the effect of the number of rungs on the probability of transition and sampling by performing SST2 simulations of the folded state of CLN025 with only 6 *λ* rungs (*F*_6 *rungs*_). This subset showed a reduced overlap in the acceptance ratio energy (Fig.S2B), resulting in a transition probability of about 44% (Fig. S2A). Remarkably, for CLN025 the SST2 simulations achieved a similar order of transition probability as the 20 temperature rungs of ST simulations, despite using only 6 *λ* rungs.

With 10 *λ* rungs, REST2 simulations showed a lower acceptance ratio of 32% and 23% for CLN025 and Trp-Cage, respectively. REST2 simulations showed a more than twofold lower acceptance ratio, which was expected as two consecutive replicas must be compatible to be swapped. For SST1 simulations, the acceptance ratio was in the same order of magnitude as for SST2 simulations with 75% and 58% for CLN025 and Trp-Cage, respectively.

#### Dependance of folding state with the *λ*

##### value

The co-evolution of RMSD and *λ* values during a subset of CLN025 *F* (1) (the number between parentheses indicates the replicas number) simulation is shown in Fig. 2. The RMSD of the backbone atoms of CLN025 to the 5awl structure oscillated between values around 0.1 *nm* in the folded state and 0.5 *nm* in the unfolded state (Fig. 2A). This RMSD was correlated with the *λ* value of the SST2 simulation; as the temperature increased or *λ* decreased, the RMSD of CLN025 increased (Fig. 2). The correlation between CLN025 RMSD and *λ* value was consistent for other SST2, SST1, ST (Fig. S5 and S6) and REST2 simulations (Fig. S7 and S8). The same observation could be made for Trp-Cage (Fig. S11, S10, S9 and S12). As ST, SST1 and REST2, SST2 simulations showed a correlation between the folding state of the protein and the temperature or *λ* value of the ladder.

**Figure 2.**
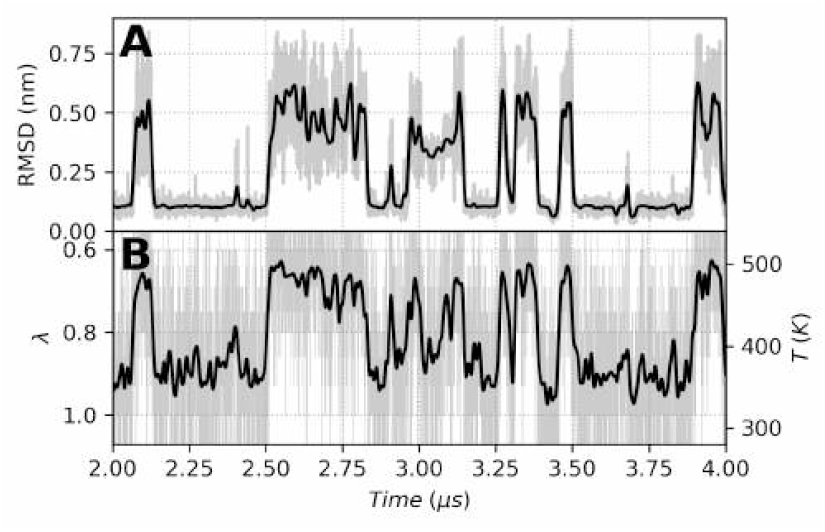
Coevolution of RMSD and *λ* value during a subset of simulation CLN025 F(1). **(A)** RMSD of backbone atoms of chignolin to 5awl structure. **(B)** *λ* evolution as a function of time, the equivalent solute-solute corresponding temperature is indicated of right y axis. To note the *λ* axis is inverted to highlight the correlation between RMSD and *λ*. For clarity, only a 2.0 *µs* subset of *F* (1) simulation out of 10.0 *µs*is shown. For both panel, raw RMSD and *λ* values are displayed as grayline, and a smooth gaussian filter with *σ* = 20 applied to RMSD and *λ* values, is displayed as black line.

#### Effect of Proline *cis-trans* transition on sampling

Proline residues occasionally undergo *cis-trans* transitions of the *ω* dihedral angle, which can significantly influence protein conformational sampling. In some simulations, such as CLN025 *F*_6;*rungs*_(1), the transition to the *cis* state trapped the protein in an unfolded conformation for several microseconds (Fig. 3). This illustrates how rare events involving proline isomerization may strongly bias folding dynamics. To avoid such artifacts, we excluded simulation frames containing prolines in the *cis* state from our comparative efficiency analyses. We further verified that this issue can be effectively addressed either by excluding *ω* dihedral angles from the solute-scaled intramolecular energy term, thereby preventing the transition, or by raising the maximum temperature (e.g., up to 700 *K*) to accelerate isomerization. Both strategies preserve the reliability of fold-unfold sampling in SST2.

**Figure 3.**
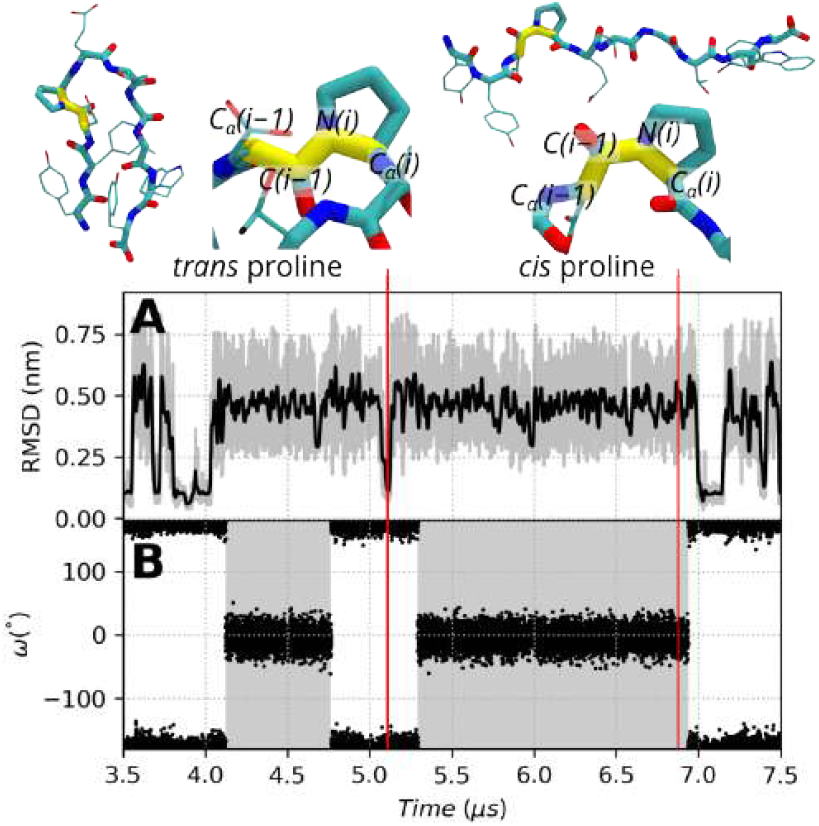
Coevolution of RMSD and Proline 4 *ω* dihedral angle during a subset of simulation CLN025. *F*_6 *rungs*_(1). (A) RMSD of backbone atoms of chignolin to 5awl structure. Raw RMSD values are displayed as gray line, and a smooth gaussian filter with *σ* = 20 applied to RMSD and *λ* values, is displayed as black line. (B) Proline 4 *ω* dihedral angle as function of time. A gray background is used to highlight when the *ω* dihedral angle is in the *cis* state. *ω* angle is defined as the dihedral angle between the C_*α*(*i−*1)_-C_(*i−*1)_-N_(*i*)_-C_*α*(*i*)_ atoms, where *i* is the Proline residue index. For clarity, only a 4.0 *µs* subset of *F*_6 *rungs*_(1) simulation out of 10.0 *µs* is shown.

Further details, including additional TrpCage and high-temperature simulations, are provided in the Supplementary Information.

#### Fold/Unfold transitions

RMSD oscillation of CLN025 and Trp-Cage appears to be of same order for ST, SST1, SST2 and REST2 simulations, when starting from a folded state or an unfolded state (Fig. S6, S10, S8 and S12). To evaluate the sampling efficiency of the different algorithms, we computed the number of transitions between folded and unfolded states per *µs* for each simulation algorithm. Chignolin and Trp-Cage were considered folded when the RMSD of the backbone atoms to the 5awl and 1l2y structures, respectively, was below 0.2 *nm* (as in Ref.^18^) and unfolded when the RMSD was above 0.25 *nm*. The number of fold/unfold transitions per *µs* for CLN025 is shown in Fig. 4A. SST2 and REST2 simulations were of the same order (about 25 transitions per computed *µs*), with a slight advantage for REST2 simulations, as ST and SST1 simulations showed slightly fewer transitions (about 15–20 transitions per computed *µs*).

**Figure 4.**
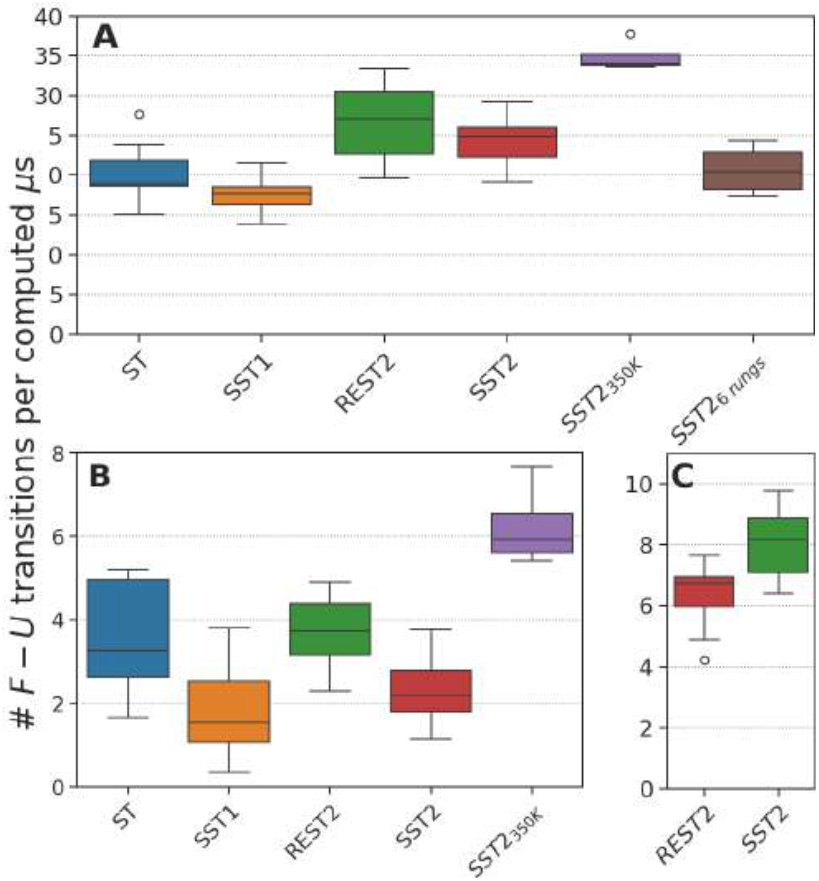
Fold/Unfold or bind/unbound transitions. Number of fold/unfold transition per *µs* for CLN025 (A), Trp-Cage simulations (B) and number of bind/unbound transitions for p97/PNGase Complex (C). Simulation frames with Proline *ω* angle in *cis* conformation have been excluded to compute fraction of folded conformation. Boxes represent the quartiles, while the whiskers show the rest of the distribution computed over the 4 to 8 replicas. ST, SST1, REST2, SST2, *SST* 2_350*K*_ and *SST* 2_6 *rungs*_ simulations are represented as blue, orange, green, red, violet and brown bars, respectively.

To understand the effect of the reference temperature *T*_*ref*_ value on sampling efficiency, we performed SST2 simulations of CLN025 starting from a folded state with *T*_*ref*_ = 350 *K* (*F*_350*K*_). As shown in Fig. 4A, the number of transitions per *µs* was significantly larger than all other simulations, averaging about 35 transitions per computed *µs*.

For Trp-Cage, the number of transitions between the folded and unfolded states seems to be an order of magnitude lower than in CLN025 (Fig. 4B). ST, SST1 SST2 and REST2 simulations were in the same order of magnitude (about 2 to 4 transitions per computed *µs*). Differences were observed between, on the one hand, SST1 and SST2 simulations, with about 2 transitions per computed *µs* (1.8 ± 1.2 and 2.3 ± 1.0 transitions per computed *µs*, respectively), and, on the other hand, ST and REST2 simulations with about 4 transitions per computed *µs* (3.5 ± 1.3 and 3.7 ± 0.9 transitions per computed *µs*, respectively). Note that the number of transitions per *µs* was higher for SST2 simulations than for SST1 simulations.

On the other hand changing the reference temperature to 350 *K* (*F*_350*K*_) resulted in a higher number of transitions (6.2 ± 1.0 transitions per computed *µs*) than all other simulations. The difference between *T*_*ref*_ = 350 *K* and *T*_*ref*_ = 300 *K* was more prononced for the Trp-Cage simulations than for the CLN025 simulations, with a difference of more than a factor 2.

#### Melting temperature

We computed the fraction of folded conformation as function of temperature or *λ* for each simulation. For REST2 and SST2 simulations we computed the effective temperature using the equation (18) proposed by Stirnemann and Sterpone.^18^

The stability curve of CLN025, depicting the fraction of folded conformation as a function of temperature, is shown in Fig. 5A. The melting temperature (*T*_*m*_) of CLN025 was estimated as 343 ± 4*K* for *SST* 2, 335 ± 11 *K* for *REST* 2, in close agreement with the experimental melting temperature of 343 *K*.^25^ For ST simulations, the melting temperature was 385 ± 13 *K*. REST2 and SST2 simulations accurately estimated the melting temperature of CLN025, whereas Stirnemann and Sterpone ^18^ underestimated the melting temperature by 20 *K* with REST2 but with a different force field.

**Figure 5.**
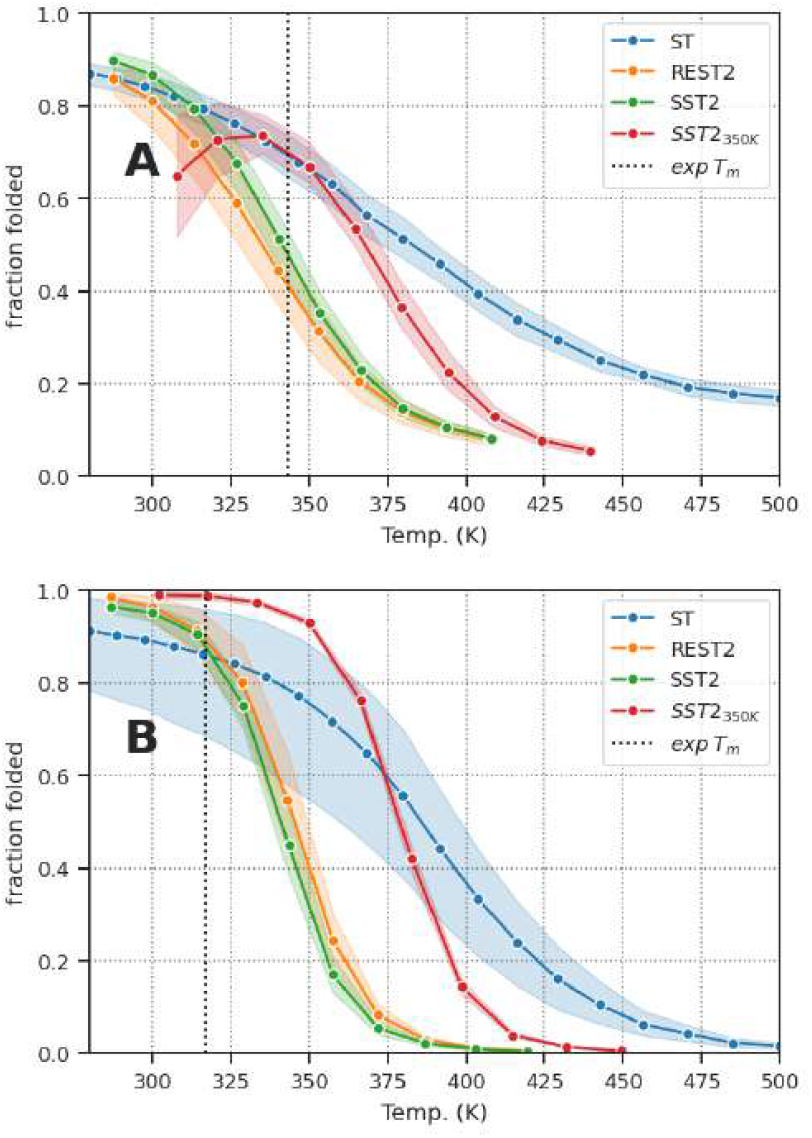
Stability Curve. Fraction of folded conformation as function of temperature for CLN025 (A) and Trp-Cage simulations (B). Simulation frames with Proline *ω* angle in *cis* conformation have been excluded to compute fraction of folded conformation. Transparent layer around the plain line displays the 95 % confidence interval computed over the 4 replicas. CLN025 and Trp-Cage were considered folded when their RMSD was below 0.2 *nm*.

However, increasing the reference temperature to 350 *K* (*SST* 2_350*K*_) resulted in a higher melting temperature of 358 ± 11*K*, overestimating the experimental value by 20 *K*.This overestimation was of the same order of magnitude as in the SST1 simulation (363.4 ± 17 *K*, where the melting temperature was computed without temperature correction as in SST2, see Fig. S29).

Fig. 5B shows the stability curve of Trp-Cage, plotting the fraction of folded conformation as a function of temperature. The melting temperature (*T*_*m*_) of Trp-Cage was estimated to be 341 ± 5, and 346 ± 4 *K* for *SST* 2 and *REST* 2, respectively, with a ~20 *K* overestimation of the experimental melting temperature of 317 *K*.^27^

For *SST* 2_350*K*_ the overestimation was even more pronounced at 379 ± 1 *K*. This overestimation was of the same order of magnitude as in the ST simulations (378 ± 42 *K*), as SST1 simulations showed a reduced overestimation with a melting temperature of 358 ± 23 *K*, (see Fig. S30 for details).

Compared to Stirnemann’s REST2 simulations which predict a melting temperature of about 320 *K*,^18^ our simulations overestimate the melting temperature by 20 *K*. In the Stirnemann’s REST2 simulations, the force field used was Amberff99SB,^26^ while in our simulations we used Amberff14SB. ^32^ This difference in force field could explain the difference in melting temperature for CLN025 and TrpCage, especially as the force field has been shown to have a strong influence on the stability of the Trp-cage.^27^

A large error bar was observed for Trp-Cage *ST* simulations. The main reason seems to be that during the *F*_*ST*_ (4) simulation, the protein was trapped in an unfolded state for more than 82 % of the simulation time. The proline 18 was in the *cis* conformation for more than 32 *µs*. As we excluded frame with proline *cis* conformation, the melting curve in simulation *F*_*ST*_ (4) was computed on only *~* 7 *µs* of the simulation time. During the first 33 *µs* of the *F*_*ST*_ (4) simulation the protein was reaching the folded state in only 3.6 % of the simulation time, as for all ST simulations the folded state was reached in 61.1% of the simulation time. This low frequency of folded state may have strongly affected the weight estimation of the *F*_*ST*_ (4) simulation, and consequently biaising the stability curve as function of temperature. This might explain the high divergence from the other simulations (Fig. S29).

Our results show that SST2 simulations provide an efficient alternative to ST simulations for sampling protein folding as a function of temperature, achieving comparable sampling efficiency with fewer rungs. In the two *toy systems*, SST2 simulations show more transitions per *µs* than SST1 simulations. Comparison of REST2 and SST2 simulation was different for the two systems. For chignolin, the number of transitions per computed *µs* was similar for SST2 and REST2 simulations, but for Trp-Cage the number of transition was than twofold lower for SST2 compare to REST2 simulations.

The correlation between RMSD and *λ* values provides insight into the conformational dynamics of the studied proteins. The accurate estimation of melting temperatures suggests the applicability of SST2, as REST2 simulations for studying protein folding as a function of temperature at a moderate computational cost (in the order of 500 GPU hours, using a nonprofessional GPU card, with 2 CPUs), but with an easier computation implementation for SST2 than REST2. SST2 is as easy as launching a standart openmm simulation, as REST2 parallelisation can be more complex to implement efficiently.

#### p97/PNGase Complex

For a protein-peptide complex, SST2 and REST2 ensured effective sampling of binding and unbinding events (Fig. S24, S25 and 6C), with an average of 8.0 ± 1.1 and 6.3 ± 3.9 binding events per computed *µs* for SST2 and REST2, respectively (peptide was considered bound if its backbone RMSD to 2hpl was less than 0.5*nm*, as it was considered unbound if its RMSD was greater than 1.0*nm*). However, it was observed that in several cases, the peptide was located in the binding site in a conformation that was not considered to be bound (with RMSD to the bound conformation 2hpl ^31^ between 0.5 *nm* and 1.0 *nm*).

We computed the PCA on SST2 and REST2 aggregated simulations, on the peptide backbone atom coordinates and used the first four components to cluster the binding sites using HDBSCAN. Analysing all the aggregated data (SST2 and REST2 simultions), the largest cluster, is the closest to the bound conformation, with an average RMSD of 0.44 ± 0.32 *nm* to 2hpl and contains the largest fraction of the simulation frames (63.5%), the second largest cluster contains only 3.0 % of the simulation frames. The largest cluster consisted of 97 % of the simulation frames where the peptide RMSD to 2hpl was below 1 *nm*. We later used a 1 *nm* RMSD cutoff to define the peptide conformations in the binding site.

We then analysed in detail the peptide conformations in the binding site using a PCA analysis limited to this subset of simulation frames followed by a HBSCAN clustering. The clustering results revealed five distinct conformations of the peptide in complex with PNGase (Fig. 6). Cluster 1 is the most populated conformation, encompassing 36.7 ± 10.0% of the simulation frames. With an average RMSD of 0.26 ± 0.07 *nm* to 2hpl, this cluster is the closest to the bound conformation. Clusters 2 and 5 account for 3.3 ± 3.1% and 0.6 ± 2.2% of the frames, respectively, and have an average RMSD of 0.83 ± 0.08 *nm* and 0.69 ± 0.04 *nm* respectively, suggesting conformations further away from the bound state. Cluster 3 and 4, comprising 2.2 ± 1.5% and 1.6 ± 2.7% of the frames, respectively, represents intermediate states with an average RMSD of 0.49 ± 0.06 and 0.45 ± 0.03 *nm*, respectively.

**Figure 6.**
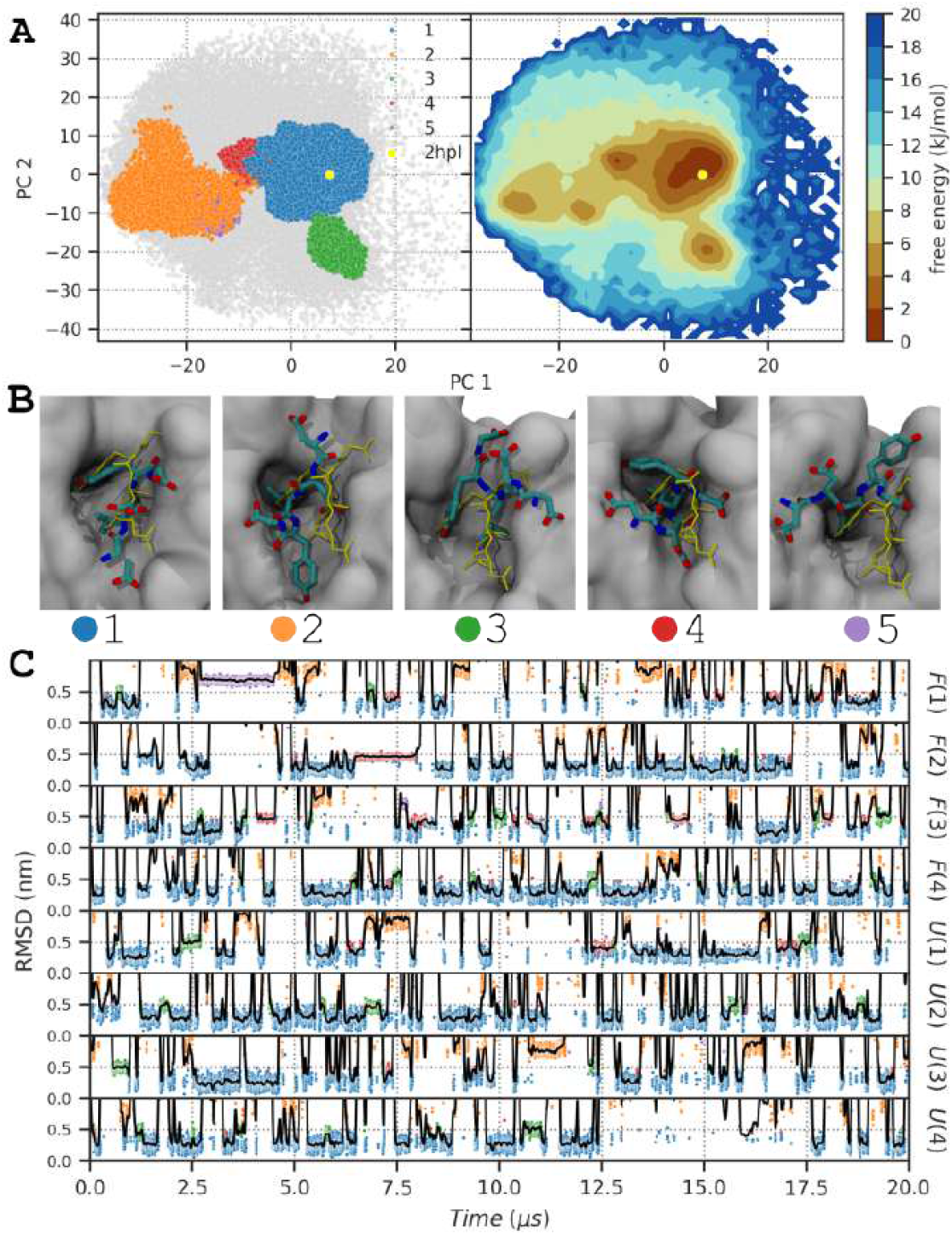
Peptide conformation in the binding site during SST2 simulations. **(A)** Projection of the two first principal component of p97 backbone atoms computed after trajectory alignement on PNGase backbone atoms. Only simulation frames where the peptide is in the binding site are displayed (peptide RMSD lower than 1.0 *nm*). Left panel displays the projection of clustered simulation frames colored by clusters, non clustered frames are displayed as gray points. Right panel displays the free energy landscape computed from the projection of the binding site simulation frames. The reference structure 2hpl ^31^ conformation is displayed as a yellow dot. **(B)** Snapshot of the five cluster structures of p97 peptide in the binding site of PNGase. Closest structure to the average structure of each cluster is displayed. PNGase is displayed as a gray surface, and the peptide heavy atoms are displayed as licorice colored by atom type. The peptide reference bound conformation (2hpl) is displayed as a yellow licorice. **(C)** RMSD as function of time for p97 peptide. RMSD has been compute on backbone atoms of p97 peptide, after structure alignement of PNGase backbone atoms, to reference structure 2hpl. Raw RMSD values are displayed as dots, colored as function of clusters, blue, orange, green, red and purple for clusters 1, 2, 3, 4 and 5, respectively. A smooth gaussian filter with *σ* = 20 applied on RMSD values, is displayed as black line. For clarity, only RMSD values between 0 and 1.0 *nm* are displayed, the complete RMSD curve is shown in the supporting information (Fig. S24).

Overall, during SST2 and REST2 simulations, the peptide explored the same conformational space (Fig. 6A and S26A), however, SST2 simulations explore the alternative conformations of the peptide in the binding site more than REST2 simulations. The occupancy of cluster 1 was 30.7 ± 10.1% and 42.8 ± 5.2% for SST2 and REST2 simulations, respectively, because alternative conformations were more populated in SST2 simulations (4.7 ± 3.7, 2.9 ± 1.5, 2.9 ± 3.5 and 1.2 ± 3.1% for clusters 2, 3, 4 and 5, respectively) than in REST2 simulations (1.8 ± 1.3, 1.5 ± 1.2, 0.3 ± 0.3 and 0.03 ± 0.07% for clusters 2, 3, 4 and 5, respectively).

In the simulation *F* (1) the peptide remained in the cluster 2 and 5 conformations for extended periods, for 11.1 and 8.8 % of the simulation time, respectively, as the residence time of cluster 1 was reduced to 17.4 % (Fig. 6).

Fig. 7 shows the fraction of peptide bound as a function of temperature for SST2 and REST2 simulations. Similarly to the fraction of folded peptide in the CLN025 and Trp-Cage simulations, the fraction of bound peptide shows a clear dependence on temperature. The curves for the *REST* 2 and *SST* 2 simulations are similar, but the error for the *SST* 2 simulations is more pronounced, mainly due to *F* (1) simulation.

**Figure 7.**
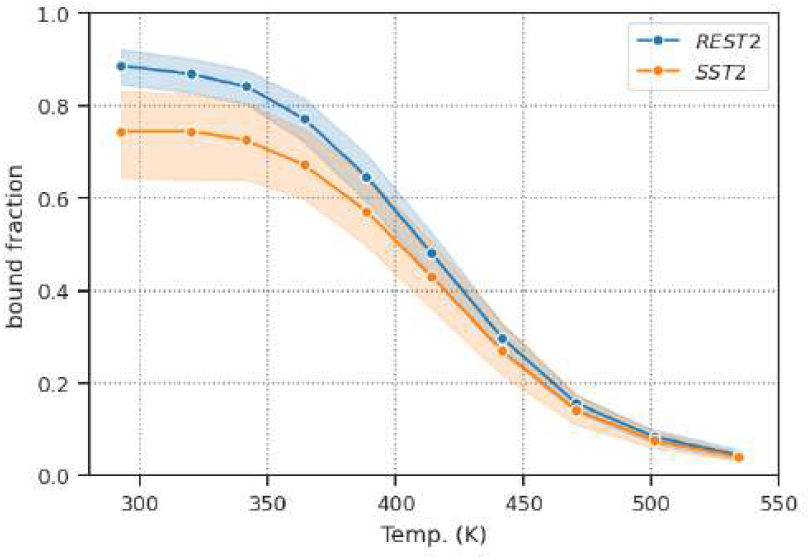
Peptide binding stability. Fraction of peptide bound as function of temperature. Peptide was considered as bound when the peptide backbone RMSD to the reference structure 2hpl^31^ was less than 0.5 *nm*. Transparent layer around the plain line displays the 95 % confidence interval computed over the 4 replicas.

During simulation *F* (1), the prolonged residence time in the intermediate conformations 2 and 5, had a marked effect on the fraction of peptide bound as a function of temperature, as this two conformations are not classified as bound. This explains the substantial error bars observed in the fraction of peptide bound as a function of temperature for the *SST* 2 simulation group (Fig. 7 and S31).

#### Simulation convergence

As shown in previous studies ^22,28,33,34^, accurate weight estimation is crucial for efficient ST simulations. The convergence of weight estimation in SST1, SST2 and ST simulations was evaluated for all systems. The results are presented in the supplementary material and demonstrate that SST1, SST2 and ST simulations exhibit comparable convergence patterns, with weight estimation reaching stability after 2–10 *µs* depending on the system. The folded or bound fraction curves as a function of temperature converge at a faster rate than the weight estimation.

Eventually, SST2 computing performance was evaluated, the computing performance was estimated to about 20 % slowdown compared to a conventional MD simulation (see Supporting Information).

### Discussion and Conclusion

The Simulated Solvent Tempering 2 (SST2) algorithm presented in this study represents a novel approach to improve the efficiency of MD simulations, particularly in the context of exploring the conformational space and thermal stability of biomolecular systems.

Our comparisons showed that SST2 achieves a performance close to ST and REST2 in terms of conformational space sampling, as in the two *toy systems*, SST2 was slightly more efficient than SST1.

We have shown that SST2 simulations can achieve comparable rung transition probabilities with less than a third of rungs required for ST simulations, and that with the same number of rungs, SST2 displayed a factor 2-3 higher probability of transtion than REST2 simulations. One must also keep in mind that, for REST2 or SST2, the number of rungs depends only on the size of the solute, it is then possible to achieve the same efficiency as ST but for larger and more complex systems. In addition, the adjustment of the reference temperature in SST2 demonstrated its potential to exceed the efficiency of ST.

One objective of our study was to assess whether SST2, similarly to REST2, could accurately reconstruct the thermal stability curves of simulated peptides. Our simulations on chignolin CLN025 and Trp-Cage showed that SST2 does indeed reproduce the thermal stability curves with comparable fidelity to REST2. It should be noted that increasing the reference temperature to 350 *K* resulted in an overestimation of the melting temperature. In fact, it is expected, since SST2 and ST simulations are consistent when *λ* = 1.0 for SST2 and when *T* = *T*_*ref*_ for ST, since they have identical Hamiltonians at these reference points. Therefore, increasing the reference temperature *T*_*ref*_ to 350*K* logically shifts the entire temperature scale upwards, resulting in a corresponding overestimation of *T*_*m*_. This behaviour, which also applies to REST2, is not due to errors in the mean-field treatment, but is a direct consequence of the choice of *T*_*ref*_. The meanfield approximation used to estimate the effective temperature estimation (eq.18) has inherent limitations. However, it is particularly useful in comparative studies, where absolute accuracy is less critical than the relative behaviour of different systems or methods. As in Timr and Sterpone,^35^ where the authors were able to use REST2 to study the stability of the chymotrypsin inhibitor in complex environments (two different protein crowded environments), and show the effect of the solvent on the stability of the protein. The demonstrated ability of SST2 to probe interactions and conformational dynamics in such complex environments suggests that it would be well suited to this type of investigation. In addition to being limited to the study of complex environments, ST and REMD are also constrained by current force fields, which are known to over-stabilise nativelike protein structures, a well-documented issue that can lead to systematic overestimation of melting temperatures across methods.

The major weakness of SST2 compared to REST2 is, as with ST compared to REMD, is the weight estimation. Stirnemann and Sterpone^18^ have shown that REST2, with 12 replicas, converges on a 150 and 250 ns time scale (corresponding to a simulated time of 1.8 and 3 *µs*) for CLN025 and Trp-Cage stability curve, respectively. In our study the SST2 simulations required 0.5-1 and 2-10 *µs* to reach convergence for CLN025 and Trp-Cage stability curves, respectively. Comparison with our REST2 simulations, shows a similar convergence for equivalent computed simulation time. SST2 achieves comparable convergence to REST2 in terms of total computation time, while also offering advantages in terms of ease of implementation. While REST2 benefits from faster convergence per unit time due to parallelism, SST2 can offer similar efficiency in terms of total computational cost, with significantly reduced technical complexity. Moreover, if a better weight estimation method was found, SST2 would theoretically be more efficient than REST2. A potential strategy to tackle this challenge is the use of machine learning-based approaches for weight estimation, which, to our knowledge, have not yet been applied to simulated tempering but could accelerate convergence and further improve the efficiency of SST2.

Another practical difference between SST2 and REST2 is that SST2 has a continuous simulation trajectory that starts from a single initial configuration. Although only the configurations at *λ* = 1.0 correspond directly to the native physical system, the uninterrupted trajectory enable a more natural dynamical evolution between exchange attempts, even when intermediate *λ* values deviate from physical reality. This continuous trajectory is particularly beneficial for capturing slow conformational or environmental relaxation processes, such as solvent rearrangements or membrane reorganization, where large-scale changes require sustained sampling rather than fragmented simulation intervals. While caution is warranted when interpreting dynamics at *λ* 1.0, the ability to track the system’s evolution across a consistent trajectory may provide insight into slow processes influencing sampling at *λ* = 1.0. For example, in the p97/PNGase complex, the RMSD of the receptor backbone over the full simulation time (0-20 and 0-1 *µs*) was slightly higher in SST2 (1.52 ± 0.40 Å) than in REST2 (1.32 ± 0.30 Å), probably reflecting the broader exploration in SST2. However, when comparing only the first microsecond (0-1 *µs*), both methods yielded similar receptor RMSD values (1.38 ± 0.28 Å for SST2 and 1.32 ± 0.30 Å for REST2). This suggests that SST2 may offer an advantage for observing slow-relaxing structural features, particularly over extended simulation times.

The accurate estimation of weights in both SST2 and ST simulations remains a significant challenge. As shown in our simulations, the weight estimation process can be affected by slow conformational transitions, and could reach convergence only after ~ 10 *µs* for the case of Trp-Cage. The utilisation of techniques, such as the Weighted Histogram Analysis Method (WHAM)^23^ has the potential to enhance the precision of weight determination, thereby addressing the inherent complexities associated with achieving reliable weight values. Nevertheless, in order to accurately estimate the weights, it is necessary to sample the folded and unfolded forms of the protein with minimum frequency at all temperatures or *λ*. This presents a challenge for proteins that fold slowly, as the simulation time must be sufficiently long to ensure adequate sampling of the conformational space.

As shown in Fig. S20 and S21, lambda rung occupation is not uniform, with low temperature rungs being more populated than others in ST and SST2 simulations. As in^28^ skewing weights towards the higher temperature rungs could improve the sampling efficiency.

One of the unexpected outcomes of the simulation analysis was the impact of proline isomerisation on the folding process. It was unexpected that, Trp-Cage could be trapped in an unfolded state for more than 32 *µs* due to a proline residue in a *cis* conformation. This observation highlights the necessity of considering slow conformational transitions in protein simulations, as they can significantly impact the accuracy of the results. This is particularly relevant, as Doshi *et. al* have shown that the free energy of *cis-trans* isomerisation is underestimated by ~6 *kcal.mol*^*−*1^ in the Amberff99SB force field.^36^ Several solutions have been proposed to address the issue of isomerisation sampling as in^37^ where the authors accelerated the sampling of N-methylated *cis-trans* amide isomers by scaling directly the force field parameters and the *ω* dihedral angle torsion constants. An alternative solution is to use a broader range of temperature for SST2, as employing temperatures =up to 700 *K* as in ^38^ accelerates significantly the time associated with proline *cistrans* transition. Nevertheless, the use of elevated temperatures implicate to use additional replicas, which could potentialy increase again the computational cost of ST and REMD simulations. Conversely, for REST2 and SST2, the number of ladders is sufficiently limited, thereby reducing the likelihood of this becoming a significant issue. The exclusion of *ω* dihedral angles from the scaled intramolecular energy term in SST2 simulations represents a significant improvement, as it allows efficient sampling of the conformational space without being hindered by slow transitions.

With the same idea, experimental data could be used to exlude additional dihedral angles from the scaled intramolecular energy, to guide or prevent the simulation to explore conformational states present or not experimentaly. While SST2 is an efficient method, it is limited by the folding or binding time of object under study. With similar concepts, derivatives of REST2 methods, such as generalized REST (gREST)^39^ restrain the scaled energy terms only to the dihedral angles and/or the CMAP terms of the solute. This enables a further reduction in the number of *λ* rungs, thereby reducing the computational cost and improving temperature sampling. This method could be adapted and implemented effortless in SST2 with the objective of enhancing the sampling efficiency of slow-folding proteins. A hybrid method combining REST2 and temperature REMD, Replica Exchange with Hybrid Tempering (REHT)^40^ have shown improvement in conformational space sampling and could be adapted to methodologies that use a single simulation, such as SST2.

Another promising direction is the integration of machine learning based post-processing approaches, such as Denoising Diffusion Probabilistic Models (DDPMs), which have recently been adapted for REMD^41^ and REST2.^42^ Since SST2 also relies on Hamiltonian scaling across modified potential energy surfaces, this concept is directly applicable. DDPMs may learn the joint distribution of solute configurations and their rescaled energies, and then exploit this knowledge to refine the reconstruction of freeenergy landscapes. Such approaches may help resolve subtle barriers that remain undersampled during the simulation itself and extend the predictive power of SST2 toward long-timescale processes without additional raw sampling.

The successful application of SST2 to chignolin CLN025 and Trp-Cage paves the way for its broader utilisation in the study of diverse biomolecular systems. Note that REST2 was clearly more efficient than SST2 for Trp-Cage simulations, with a factor 2 higher number of transitions per computed *µs*. However, as evidenced by the p97/PNGase complex simulations, SST2 is also capable of efficiently sampling large systems as protein-peptide complexes, with higher, sampling efficiency than REST2, thereby offering invaluable insights into the dynamics of ligand-receptor binding and recognition. SST2 shows great promise for applications beyond isolated small protein simulations. Its capacity to efficiently sample conformational space makes it a promising candidate for the study of systems involving ligands, peptides, or proteins within complex environments, such as lipidic membranes or in complex with a receptor. This versatility establishes SST2 as a valuable tool for investigating biomolecular interactions and dynamics in realistic settings.

In conclusion, the SST2 algorithm which is presented as an amalgamation of ST and REST2, demonstrates promising advancements in MD simulation sampling. Our comparative analysis and application to specific systems have highlighted the potential of SST2 to efficiently sample conformational space and reconstruct thermal stability curves. The adaptability of SST2, evidenced by its ability to surpass ST, SST1 and REST2 efficiency under specific conditions, positions it as a valuable addition to the toolkit of computational biophysicists.

## Materials and Methods

We first present the concepts and theories behind REMD, ST, REST, REST2, and eventually the theory and implementation of SST2.

### Classical REMD

Replica Exchange Molecular Dynamics (REMD)^9^ simulates multiple copies (or *replicas*) of a molecular system at different temperatures. Swaps between neighbouring replicas are attempted at regular time intervals. The probability of an exchange between temperature *m* and *n* is calculated using a Metropolis criterion 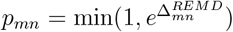 with:

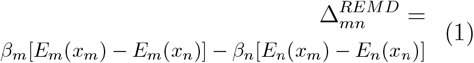

where *β*_*m*_ = ^1^*/*(*k*_*B*_*T*_*m*_), *E*_*m*_ and *x*_*m*_ are the potential energy and the configuration of the system at temperature *T*_*m*_. Since the potential energy is independent of the temperature we obtain:

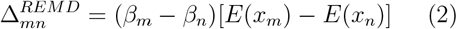

### REST

In the first version of Replica Exchange with Solute Tempering (REST),^12^ instead of just increasing the temperature as in REMD, the Hamiltonian is also updated as the temperature increases to keep the solvent at cold or ambient temperature (*T*_*ref*_) using:

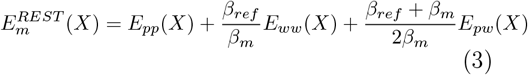

where *X* is the configuration of the system, *E*_*pp*_ is the solute intramolecular energy, *E*_*pw*_ is the solute solvent interaction energy, and *E*_*ww*_ is the solvent intramolecular energy.

Although REST has been shown to accelerate the convergence of the conformational space distribution for small molecules, it has proved less effective than temperature REMD for larger systems such as Trp-Cage and *β* hairpin.^13^ To overcome this limitation, several improvements to REST have been implemented.

Hamiltonian modifications inspired by Moors *et. al* ^43^ and Terakawa *et. al* ^44^ led to improved version of REST called REST2. In REST2^14^ simulations, the replicas are all ran at the same temperature *T*_*ref*_, but the replicas evolve with different Hamiltonians. For replica *m* running at *λ*_*m*_ = *β*_*m*_*/β*_*ref*_ = *T*_*ref*_*/T*_*m*_ (considered as an approximation of *T*_*m*_):

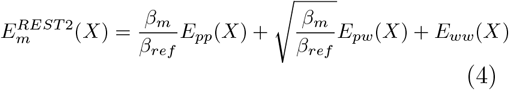

or:

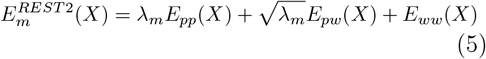

The acceptance ratio for the exchange between replicas *m* and *n* is defined by:

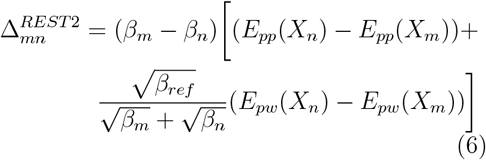

Since solute-solute interactions are scaled equally compared to solvent-solvent interactions by *β*_*m*_*/β*_*ref*_ in REST and REST2, the difference comes from the solute-solvent interaction, which is scaled by (*β* _*m*_ + *β*_*ref*_)*/*2*β*_*m*_ in REST and 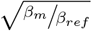 in REST2. REST2 by using a weaker solute-solvent interactions at high temperature, favours the sampling of folded conformations even at high temperatures.

### Simulated tempering

In simulated tempering (ST)^10,11^ a single copy of the system is simulated and the temperature is periodically updated between a predetermined set of values. As in REMD, the Metropolis algorithm is used to decide whether or not to accept a temperature swap, one of the main advantages over REMD is that the temperature swap depends on only one simulated system rather than two, which speeds up the temperature transition. Zhang and Ma^21^ have shown that ST gives a higher rate of traversal of energy space with the reserve of an accurate partition function (ST weights *w*_0_, *w*_1_, … *w*_*i−*1_).

The acceptance rate is given by 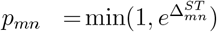 with:

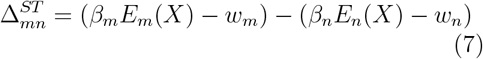

The potential energy function *E*_*m*_(*X*) is independent of the temperature *T*_*m*_, so *E*_*m*_ = *E*_*n*_, which gives:

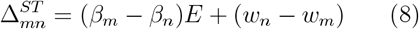

In ST simulation, the correct calculation of the exact partition function is particularly important for an efficient ST computation. To compute the weights we used the method proposed by Park and Pande:^34^

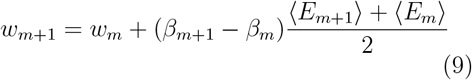

Since *E*_*m*_ must be calculated forehand, we use the *on the fly* weight calculation developed by Nguyen *et al*.^33^, which allows to approximate *w*_*m*+1_ if no simulation has been calculated at *T*_*m*+1_:

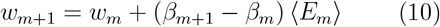

### Simulated solute tempering (SST1)

In the SST implementation of REST in ST,^23^ the authors modified the potential energy function using:

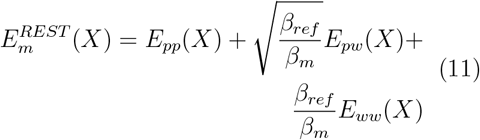

Using the equation (11) in (8) gives:

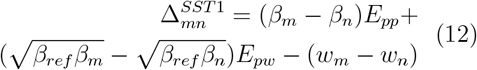

While SST1 combines the advantages of ST and REST, it suffers from the same limitations as REST and has not been adopted by the community as a reference method for protein folding.

### Simulated solute tempering 2 (SST2)

We used a slightly modified version of the original implementation of REST2. In the approach proposed in,^18^ only the proper torsion terms of the solute were scaled among the bonded terms:

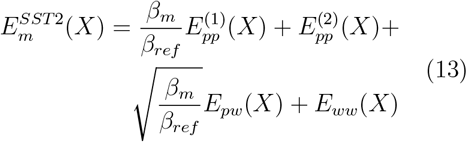

or using *λ*_*m*_ = *β*_*m*_*/β*_*ref*_ :

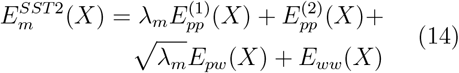

where 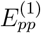 is the scaled solute intramolecular energy (LJ, Coulomb, and proper torsions), is the unscaled solute intramolecular energy (bonds, angles, and improper torsions), *E*_*pw*_ is the solute-solvent interaction energy and *E*_*ww*_ is the solvent intramolecular energy.

The acceptance ratio is given by 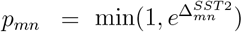, where inserting (13) into (8) gives:

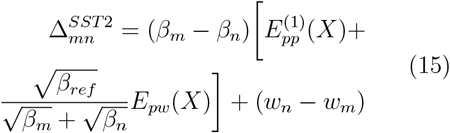

This equation is equivalent to the original SST methods (12).

Using 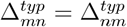, we obtain:

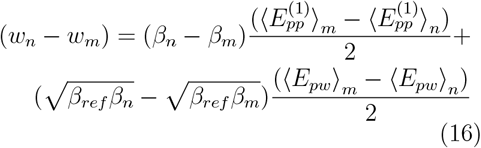

The exchange with neighboring *m* + 1 replicas is determined by the fluctuation of 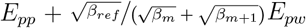 or for simplicity, since *β*_*m*_ and *β*_*m*+1_ are close, we will later monitor 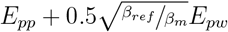.

### Temperatures distribution

*m* temperatures have been chosen to be exponentially spaced between the two extremes *T*_min_ = *T*_0_ and *T*_max_ = *T*_*m−*1_. With *k* included in [0, 1, …, *m−* 1], the temperatures have been computed as follows:

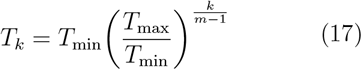

If *T*_*ref*_ is different from *T*_min_, we use the same formula to compute the effective temperatures between *T*_min_ and *T*_*ref*_ and between *T*_*ref*_ and *T*_max_.

### Effective temperature estimation

To estimate the corrected SST2 solute temperature we use the formula proposed by Stirnemann and Sterpone in:^18^

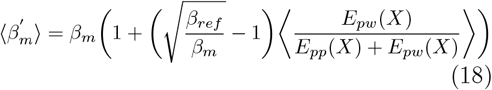

### SST2 implementation

The ST, SST1 and SST2 protocols have been implemented by Python scripts using OpenMM.^45^ ST’s original OpenMM script, written by Peter Eastman, was modified to implement the weight calculation of Park and Pande^34^ and the *on the fly* weight calculation from Nguyen *et al*.^33^ The same script was also used to write the SST2 scripts.

The scaling of non-bonded interactions was done by scaling the *ϵ* parameters of the Lennard-Jones potential by *λ*_*m*_ = *β*_*m*_*/β*_*ref*_, while the charges of solute atoms were scaled by 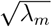. The solute intramolecular energies (bonds, angles, and improper torsions) were left unchanged, for selected proper torsion terms, the dihedral constant term *k* was scaled by *λ*_*m*_.

In order to accurately compute the different energy terms *E*_*pp*_, *E*_*pw*_ and *E*_*ww*_, two additional molecular systems were created in addition to the simulated systems, consisting of the solute atoms only, and the solvent atoms only. At each step where the energy terms had to be computed, the additional systems were assigned the same coordinates as the simulated systems. This allows the long term electrostatic contribution of *E*_*pp*_ and *E*_*ww*_ to be accurately computed and the the non-bonded interaction contribution *E*_*pw*_ to be derived.

As in classical MD simulations, it is generally recommended to neutralize the full system when using Ewald summation methods for long-range electrostatics. However, when the solute is charged, the solute-only and solvent-only subsystems are not neutral, which could in principle lead to artifacts in the energy calculation. To address this, it is important to note that PME implicitly introduces a uniform neutralizing background charge. OpenMM versions prior to 8.3.1 did not include the explicit correction term for this background, whereas from version ≥8.3.1 onward, the correction has been implemented, ensuring consistent energy reporting in charged systems. We verified that this correction has no effect on forces and only a very minor effect on reported energies. Specifically, we performed 10 ns simulations of CLN025, TrpCage, and PNGase, and compared exchange acceptance with and without the correction. In all three cases, ~99.9% of exchanges were identical (i.e., only ~5 out of 5000 attempts differed). Thus, while the impact is limited to reported energies and exchange calculations, we recommend that users employ OpenMM ≥ 8.3.1 to ensure robust and consistent treatment of charged systems.

The ST, SST1 and SST2 implementation is available at github.com/samuelmurail/SST2. To use our simulation script, we recommend using the Remd Temperature Generator webserver^46^ at https://virtualchemistry.org/remd-temperature-generator/ to calculate the SST1, SST2 or ST temperature list or simply the number of ladders. The user should divide the expected probability by 2 (ratio of expected probability exchange between REMD and ST, or REST2 and SST2). For SST2, enter the number of solute atoms for *Number of protein atoms* and 0 for *Number of water molecules*.

### System Preparation

Two types of systems were prepared, systems starting from an unfolded state, and systems starting from a folded state.

The unfolded structures were generated using the pdb manip py library (pdb-manip-py. readthedocs.io) and later the pdb numpy library (pdb-numpy.readthedocs.io) to generate a linear structure of the peptide. Structure preparation and protonation were performed using the `pdbfixer` module of the OpenMM package. The protocol for preparing the folded structure was similar to that for the unfolded structure, except that the implicit simulation step was skipped, as we started directly from the PDB structures 5AWL^29^ and 1L2Y^30^ for the chignolin, and Trp-Cage proteins, respectively.

The simulation of the p97/PNGase complex was prepared using the PDB structure 2HPL^31^ and the same protocol as for the folded structure. For the unfolded structure, the PNGase structure was prepared using the PDB structure 2HPJ,^31^ as the p97 peptide was prepared as a linear peptide and placed randomly around the PNGase. The same minimisation, equilibration and simulation protocol was used.

### Molecular Dynamics simulation preparation

The simulations were computed using the OpenMM 7.7 package.^45^ In order to circumvent the utilisation of an excessively large simulation box, the chignolin and Trp-Cage linear structures were minimised using the L-BFGS algorithm for up to 10.000 steps and then submitted to an implicit solvent equilibration of 10 *ns* using the Langevin integrator and the GBSA-OBC solvation model^47^ with the Amber99sbNMR^48^ force field. We had originally planned to use Amber14SB for implicit solvent simulations, but the implicit solvent model was not implemented in OpenMM at the beginning of the study. The obtained structures, folded structure and protein-peptide complexes, were then used as starting points for the explicit solvent simulations. The force field was Amber14SB^32^ and proteins were solvated in a truncated octahedron boxes with a padding of 1.5 *nm* and the TIP3P water model.^49^ Where necessary, counterions were added to counteract the charge of the protein. The systems were again minimised up to 10.000 steps and equilibrated for 10 *ns*, the temperature and pressure were equilibrated using a Monte Carlo barostat at a temperature of 300 *K* and a pressure of 1 atmosphere. A nonbonded cutoff of 1.0 *nm* was used with the Particle Mesh Ewald algorithm for long range electrostatic interactions. We used a 4 *fs* integration time step using heavy hydrogen assigned a mass of 3 atomic mass units.^50^

### Simulation Run

We performed ST, SST1 and SST2 simulations of CLN025 and Trp-Cage (Table 1) in explicit solvent, starting from the unfolded and folded states (*U* and *F*). Temperature updates were performed every 2 *ps* for ST, SST1 and SST2 simulations, following the protocol in, ^28^ as we used a friction coefficient of 10, 1 and 1 *ps*^*−*1^ for the Langevin integrator, for ST, SST1 and SST2 simulations, respectively (see the Supporting Information and Fig. S33 for a detailed discussion of the effect of the friction coefficient on ST and SST2 simulations). For both ST and SST2 simulations, four replicas for fold and unfold structure were computed for 10 and 40 *µs*, for CLN025 and Trp-Cage, respectively. As for SST1 simulations, we used simulation time of 10 and 20 *µs*, for CLN025 and Trp-Cage, respectively. The SST2 simulations used a reference temperature *T*_*ref*_ = 300 *K*, with 10 *λ* rungs of 1.07, 1.00, 0.93, 0.86, 0.80, 0.75, 0.69, 0.64, 0.60, and 0.56, corresponding to solute-solute corresponding temperatures of 280.0, 300.0, 322.9, 347.5, 374.0, 402.5, 433.2, 466.2, 501.7, and 540 *K* (*λ*_*m*_ = *T*_*ref*_*/T*_*m*_). We used one temperature of 280 *K* and 9 discrete temperatures exponentially distributed from 300 to 540 *K*.

For comparison, ST simulations of CLN025 and Trp-Cage in unfolded and folded initial conformations (*U*_*ST*_ and *F*_*ST*_) were performed with 20 temperature rungs ranging from 280.0 to 500.0 *K*. As shown in,^18^ the solute-solute temperature in SST2 simulations is not the effective temperature of the solute, as the solvent is being kept at a reference temperature and the solute-solvent interactions being scaled by 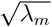. The effective solute temperature results from a balance between the two scaled energies *E*_*pp*_ and *E*_*pw*_. Temperature ranges in ST and SST2 were chosen to approximate the two exteme temperatures in the stability curve for CLN025 and Trp-Cage (folded fraction around 1.0 and 0.0), this explain the difference in the range of temperature between ST and SST2 simulations.

The effect of changing the reference temperature to a higher value was investigated for CLN025 and Trp-Cage. SST2 simulations of CLN025 and Trp-Cage in folded states were performed with 10 *λ* rungs, ranging from 280.0 to 540.0 *K* with *T*_*ref*_ = 350*K*, resulting in *λ* values of 1.25, 1.16, 1.08, 1.0, 0.93, 0.87, 0.81, 0.75, 0.70, and 0.65, corresponding to solute-solute equivalent temperatures of 280.0, 301.6, 324.9, 350, 376.2, 404.4, 434.7, 467.3, 502.3, and 540 *K*. Note that if the solute-solute corresponding temperature distribution is similar at *T*_*ref*_ = 300*K* and *T*_*ref*_ = 350*K*, the temperature of the solvent is simulated at the reference temperature, resulting in a 50*K* temperature difference.

Given the size of a protein-peptide complexes, during a classical ST simulation, we would need to use an excessive number of ladders to ensure a good exchange rate. Consequently, as the temperature rises, the receptor is likely to unfold, and adding position restraints to maintain the receptor structure could be required. For this reason, we chose to use only the SST2 algorithm to simulate the p97/PNGase complex. In these simulations, only the peptide ligand was considered as part of the solute subject to scaling, while the receptor remained unscaled. Importantly, no constraints were applied to either the receptor or the ligand, allowing the system to evolve freely under the SST2 framework.

To assess SST2’s performance in simulating protein-peptide interactions, we performed eight molecular dynamics (MD) simulations of the p97/PNGase complex of 20 *µs*, four starting from the folded state and four starting from the unfolded state in explicit solvent (Table 1). To increase the sampling efficiency, we use a reference temperature *Tref* = 320*K*, resulting in 10 *λ* rungs of 1.14, 1.00, 0.91, 0.82, 0.75, 0.68, 0.07, 0.56, 0.50, 0.46, corresponding to solute-solute corresponding temperatures of 280.0, 320.0, 352.9, 389.2, 429.2, 473.3, 521.9, 575.6, 634.8, and 700 *K*. Preliminary tests indicated that a maximum solute-solute temperature of 700 *K* was optimal for the system, allowing us to obtain a corrected maximum temperature of 500 *K*. At this temperature, the binding percentage was observed to be close to 0%, providing an ideal condition for evaluating the performance of the algorithm.

#### Exclusion of proline *ω* dihedral angles

As will be shown later, CLN025 proline 4 in the *cis* conformation can trap the protein in an unfolded state. To avoid this issue, an additional option was added to the SST2 script, to exclude proline *ω* dihedral angles from the solute scaled intramolecular energy term 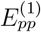 and keep them in the unscaled solute intramolecular energy term 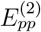. To do this, all dihedral terms containing atoms N_(*i*)_ and C_(*i−*1)_ (where (*i*) indicates the proline residue number) at positons 2 and 3 of the dihedral term were excluded from 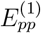 and kept in 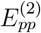. This option was enabled on replicas 3 and 4 of Trp-Cage *F* and *U* simulations, and on all replicas of Trp-Cage *F*_350*K*_ simulations (Table 1).

### REST2 simulations

To conduct REST2 simulations we used the Psivant implementation of REST2 in openmm, the femto package was taken from `github. com/Psivant/femto.` Same protocol for system preparation was used as in SST2, a 2 *ps* interval was used to attempt lambda swap between neighboring lambdas. 10 lambdas were used with the same temperatures as used in SST2 simulations. For CLN025, 4 replicas of 1.0 *µs* were ran starting from folded and unfolded forms, for TrpCage, 4 replicas of 2.0 *µs* were ran starting from folded and unfolded forms, as for p97/PNGase complex (only the peptide ligand is considered as solute), 4 replicas of 1.0 *µs* were ran starting from bounded and unbounded forms (Table 1). The first 10% of the simulations were removed for analysis.

### Analysis

Structural alignment and Root Mean Square Deviation (RMSD) were computed on the backbone atoms by removing the first and last residues of chignolin (residues 2 to 9 were used) and the two first and two last residues of TrpCage (residues 3 to 18 were used). In the case of the p97/PNGase complex, trajectory alignment was performed on the PNGase backbone atoms, and the RMSD was computed using all backbone atoms of p97.

In order to gain a detailed inderstanding of the p97 peptide conformation within the PNGase binding site, a Principal Component Analysis (PCA) was conducted on simulation frames with a peptide RMSD of less than 1.0 *nm*. Subsequently, the resulting peptide conformations were clustered using the HDBSCAN algorithm^51^ on the first four components.

The `MDAnalysis`^52^ python package was used together with some home-made python scripts to analyse the MD simulations and perform statistical analysis and plotting.

The python package `scipy`^53^ was used to compute four parameter logistic (4PL) regression on folding fraction as function of temperature curves to extract the melting temperature. The equation model used is:

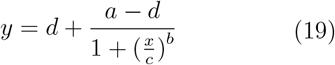

where *a* is the folded fraction at low temperature, *d* is the folded fraction at high temperature, *c* is the inflection point, also called the melting temperature *T*_*m*_, and *b* is the Hill’s slope of the curve (or the steepness of the curve at *T*_*m*_).

## Supporting information

Supplementary Informations

## Acknowledgement

The authors would like to express their gratitude to Fabio Sterpone and Samuela Pasquali for their valuable contribution to the discussion surrounding the theoretical aspects and implementation of SST2. Furthermore, we extend our appreciation to Peter Eastman and more broadly, to the entire OpenMM development team, for their efforts in making OpenMM and the simulated tempering implementation accessible as an open-source resource.

## Author contributions

S.M. did the method development and implementation, MD simulations preparation and run. D.S., G.M., P.T. and S.M. analyse the simulations and wrote the paper.

## Code availability

The various Python scripts used in this work are available on github at `github. com/samuelmurail/SST2.` Simulation trajectories have been deposited at zenodo.org/records/13772542 with the doi 10.5281/zenodo.13772541.

## Supporting Information Available

Details about the simulations, discussion of temperature swap interval, analysis about SST2 computing performance, ST, SST1, REST2 and SST2 simulations convergence.

