## Supplementary Informations for "Simulated Solute Tempering 2: An Efficient and Practical Approach to Protein Conformational Sampling and Binding Events"

### 1 CLN025 and Trp-Cage SST2 simulations

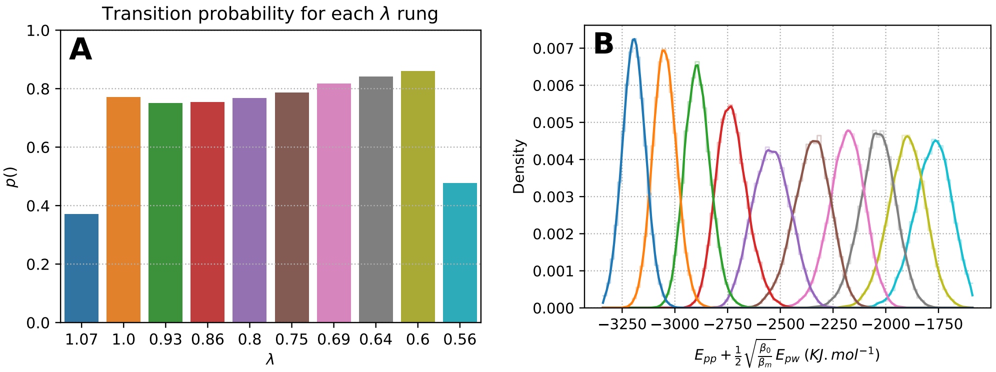


Figure S1: **Probability of transition and distribution of the SST2 energy as function of $\lambda_i$ for simulation Trp-Cage $F$ (1). (A)** Probability of transition at each $\lambda$ rung. **(B)** Distribution of acceptance ratio energy $E_{pp} + 0.5\sqrt{\beta_{ref}/\beta_m}E_{pw}$ for each $\lambda$, the color code for $\lambda$ value is identical to the upper panel one. Maximum $\lambda$ value 1.07 corresponds to a solute-solute corresponding temperature of 280 $K$, as minimum $\lambda$ value 0.56, correspond to a temperature of 540 $K$.

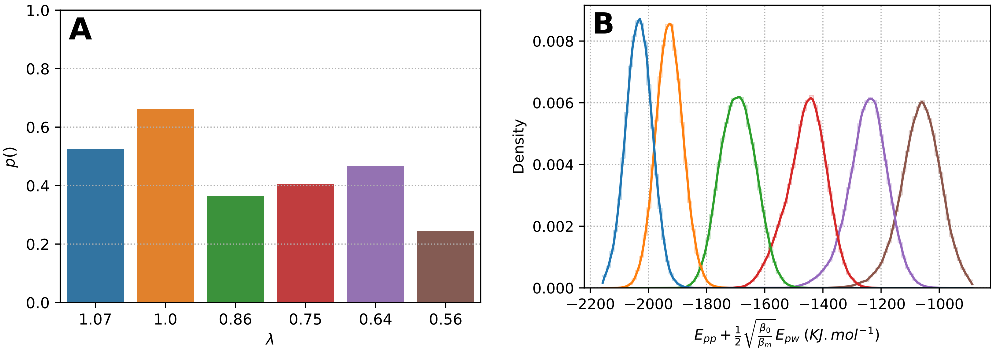


Figure S2: **Probability of transition and distribution of the SST2 energy as function of $\lambda_i$ for simulation CLN025 $F_{6rungs}$ (1). (A)** Probability of transition at each $\lambda$ rung. **(B)** Distribution of acceptance ratio energy $E_{pp} + 0.5\sqrt{\beta_{ref}/\beta_m}E_{pw}$ for each $\lambda$, the color code for $\lambda$ value is identical to the upper panel one. Maximum $\lambda$ value 1.07 corresponds to a solute-solute corresponding temperature of 280 $K$, as minimum $\lambda$ value 0.56, correspond to a temperature of 540 $K$.

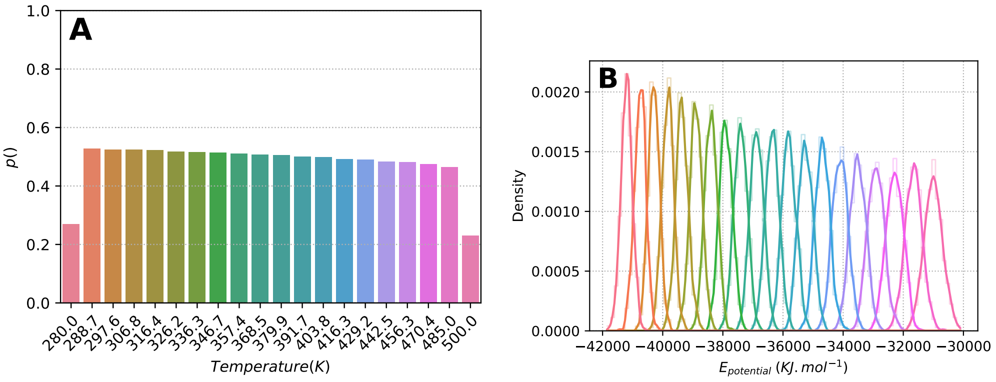


Figure S3: **Probability of transition and distribution of the potential energy as function of temperature for simulation CLN025 $F_{ST}$ (1). (A)** Probability of transition at each temperature rung. **(B)** Distribution of acceptance ratio energy $E_p$ for each temperature, the color code for temperature value is identical to the upper panel one.

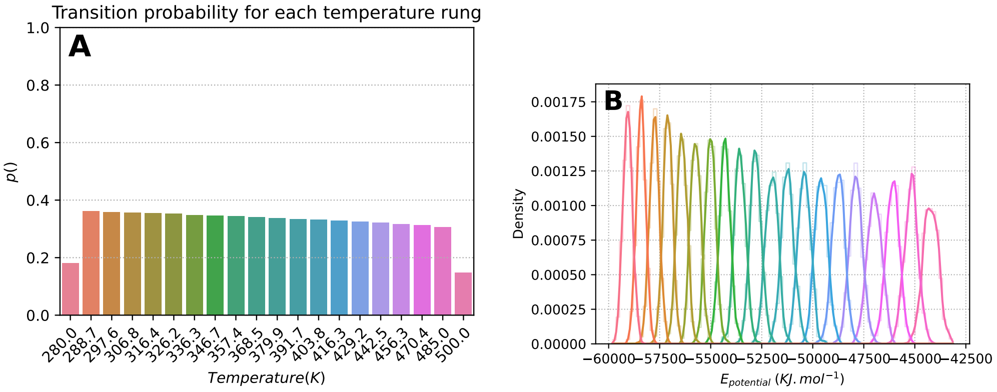


Figure S4: **Probability of transition and distribution of the potential energy as function of temperature for simulation Trp-Cage $F_{ST}$ (1). (A)** Probability of transition at each temperature rung. **(B)** Distribution of acceptance ratio energy $E_p$ for each temperature, the color code for temperature value is identical to the upper panel one.

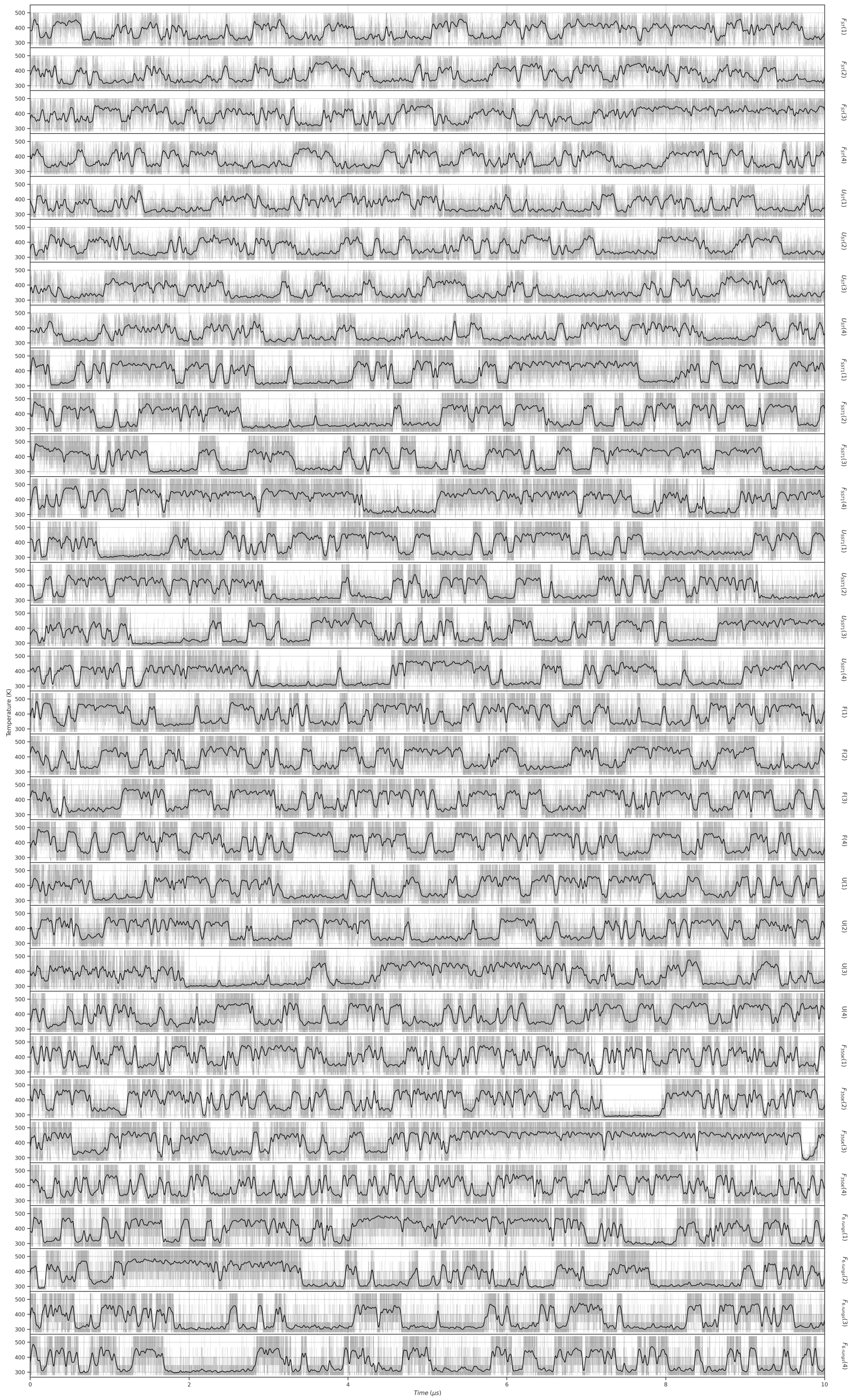


Figure S5: **Temperature as function of time for CLN025 simulations.** For SST2 simulations, the temperature correspond to the solute-solute corresponding temperature. Raw temperature values are displayed as gray line, and a smooth gaussian filter with $\sigma = 20$ is applied, and displayed as black line.

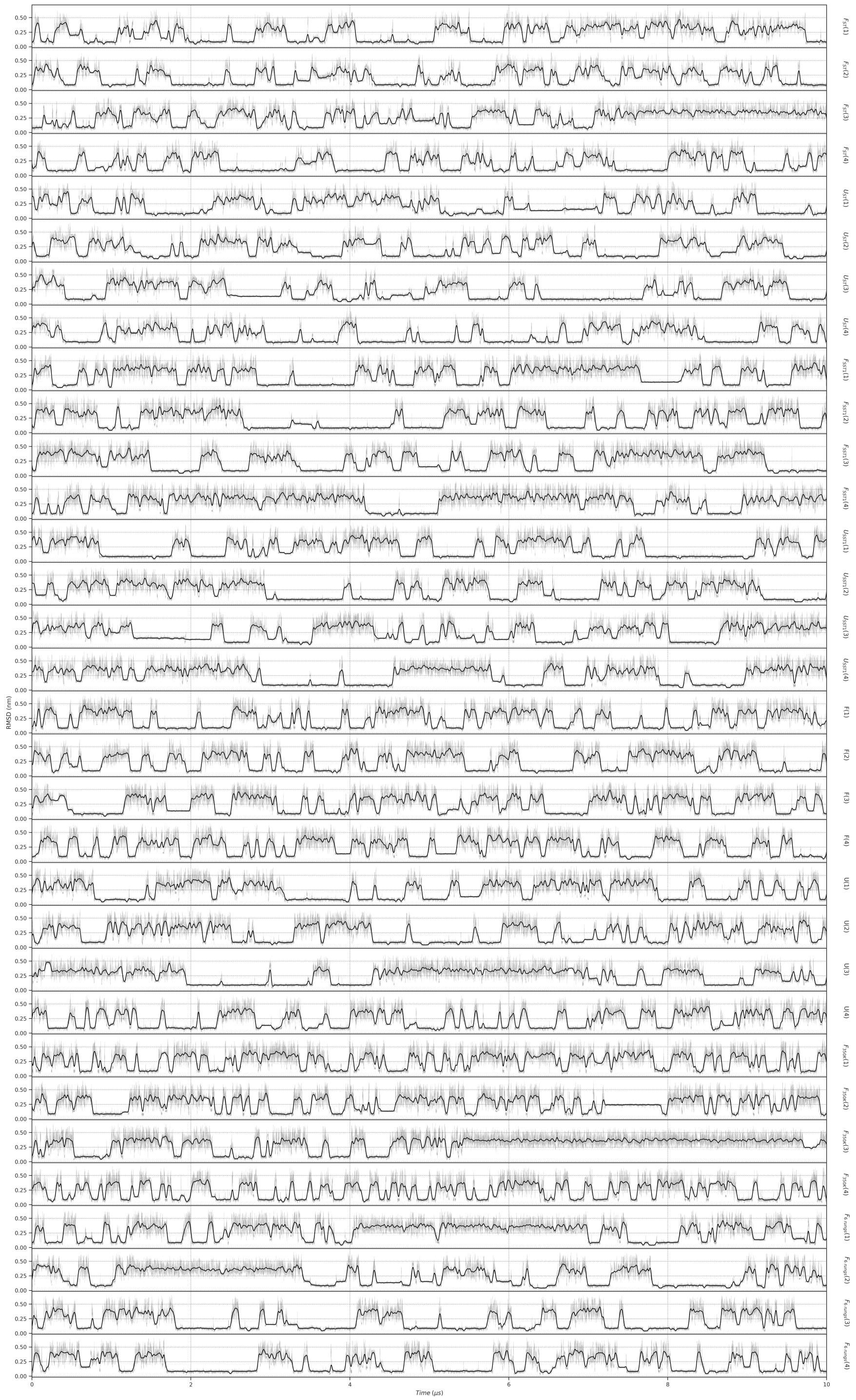


Figure S6: **RMSD as function of time for CLN025 simulations.** RMSD has been compute on backbone atoms, without the first and last residues, to reference structure 5awl.[1] Raw RMSD values are displayed as gray line, and a smooth gaussian filter with $\sigma = 20$ is applied, and displayed as black line.

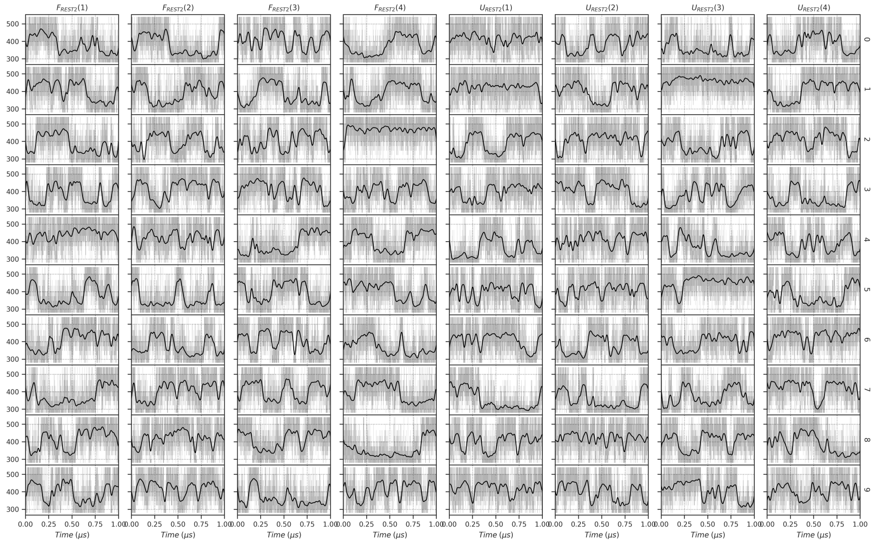


Figure S7: **Temperature as function of time for REST2 CLN025 simulations.** The temperature correspond to the solute-solute corresponding temperature. Raw temperature values are displayed as gray line, and a smooth gaussian filter with $\sigma = 20$ is applied, and displayed as black line.

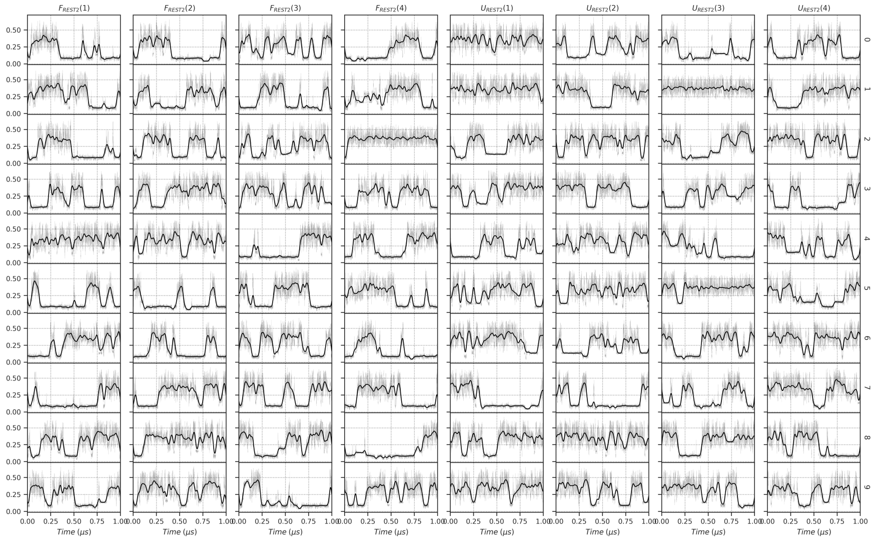


Figure S8: **RMSD as function of time for REST2 CLN025 simulations.** RMSD has been compute on backbone atoms, without the first and last residues, to reference structure 5awl.[1] Raw RMSD values are displayed as gray line, and a smooth gaussian filter with $\sigma = 20$ is applied, and displayed as black line.

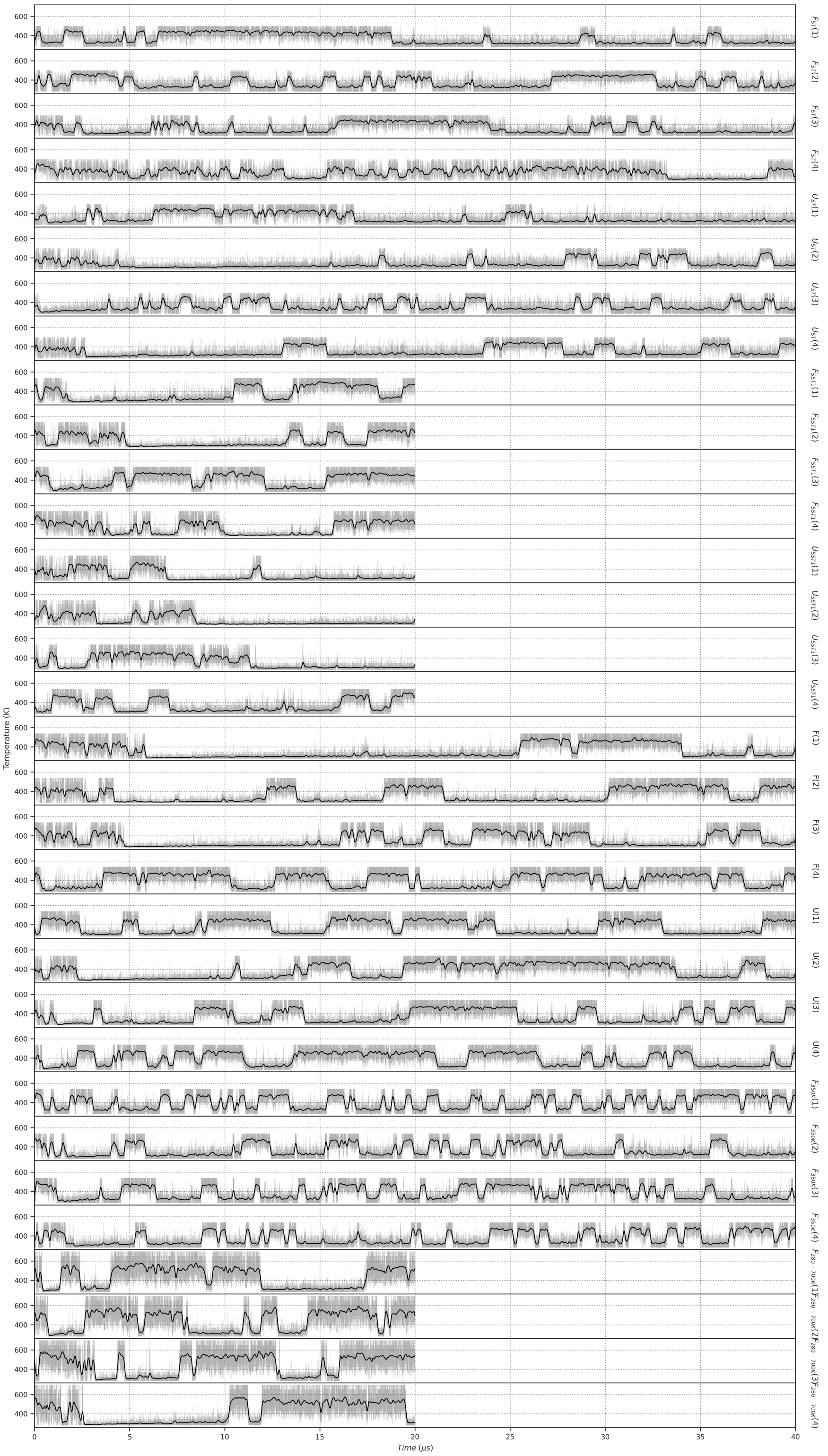


Figure S9: **Temperature as function of time for Trp-Cage simulations.** For SST2 simulations, the temperature correspond to the solute-solute corresponding temperature. Raw temperature values are displayed as gray line, and a smooth gaussian filter with $\sigma = 20$ is applied, and displayed as black line.

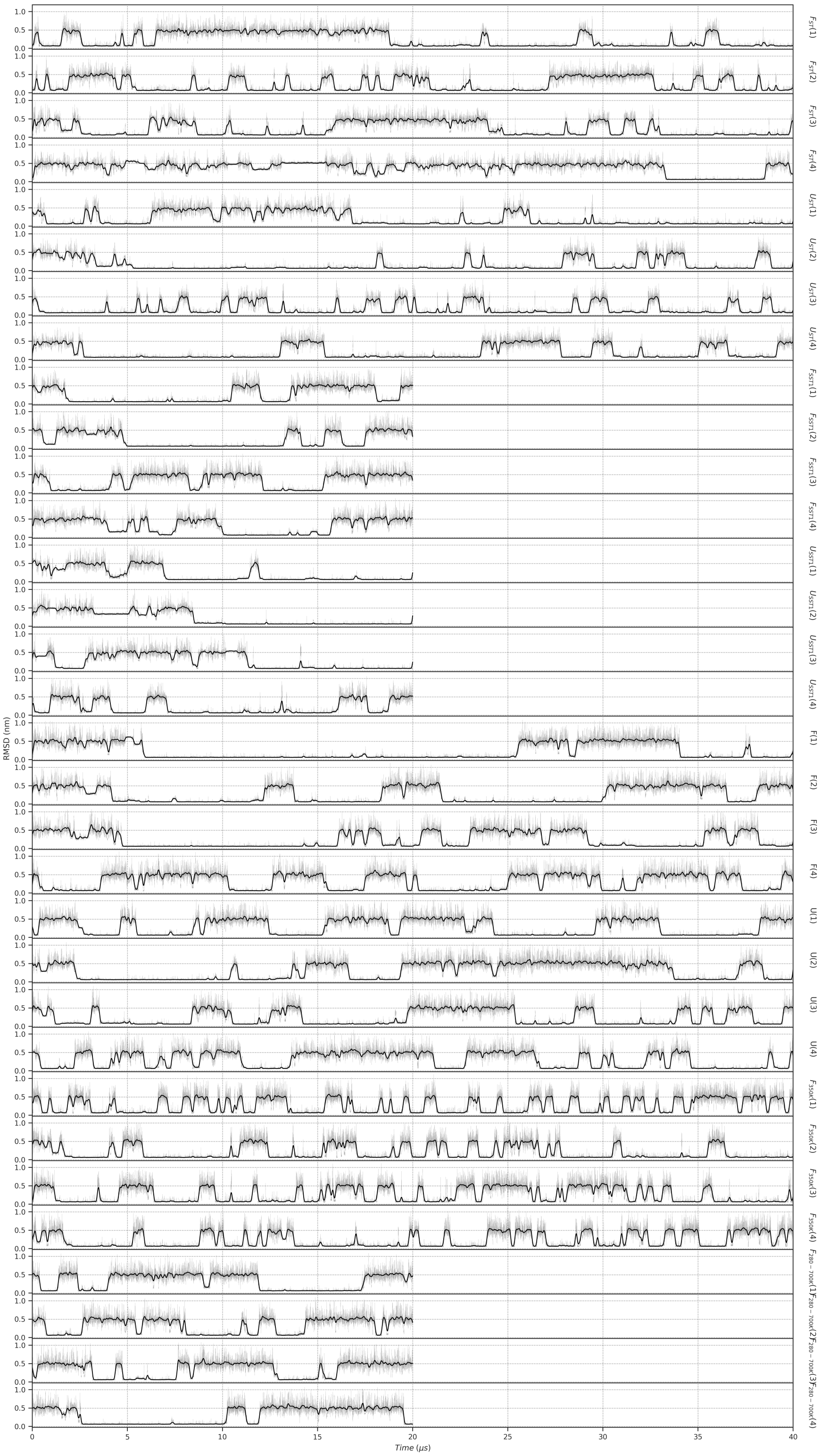

Figure S10: **RMSD as function of time for Trp-Cage simulations.** RMSD has been compute on backbone atoms, without the first and last 2 residues, to reference structure 1l2y.[2] Raw RMSD values are displayed as gray line, and a smooth gaussian filter with $\sigma = 20$ is applied, and displayed as black line.

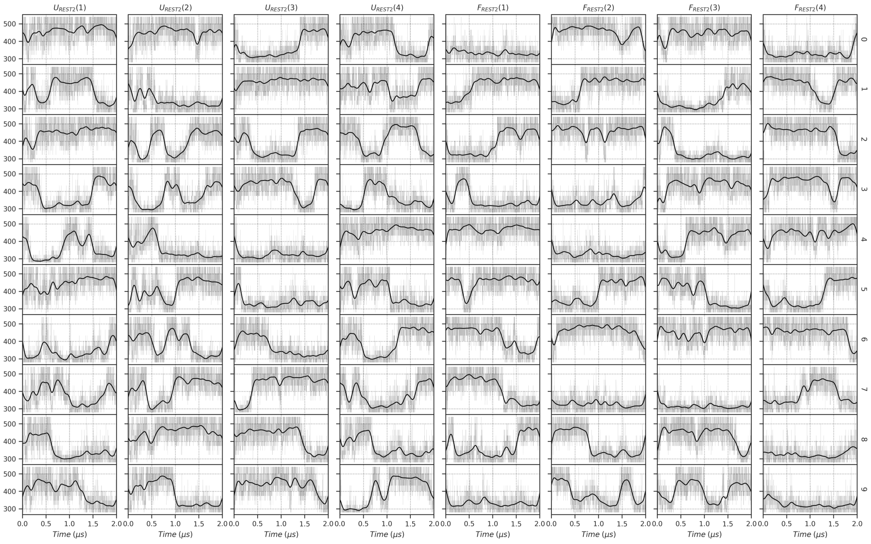

Figure S11: **Temperature as function of time for REST2 TrpCage simulations.** The temperature correspond to the solute-solute corresponding temperature. Raw temperature values are displayed as gray line, and a smooth gaussian filter with $\sigma = 20$ is applied, and displayed as black line.

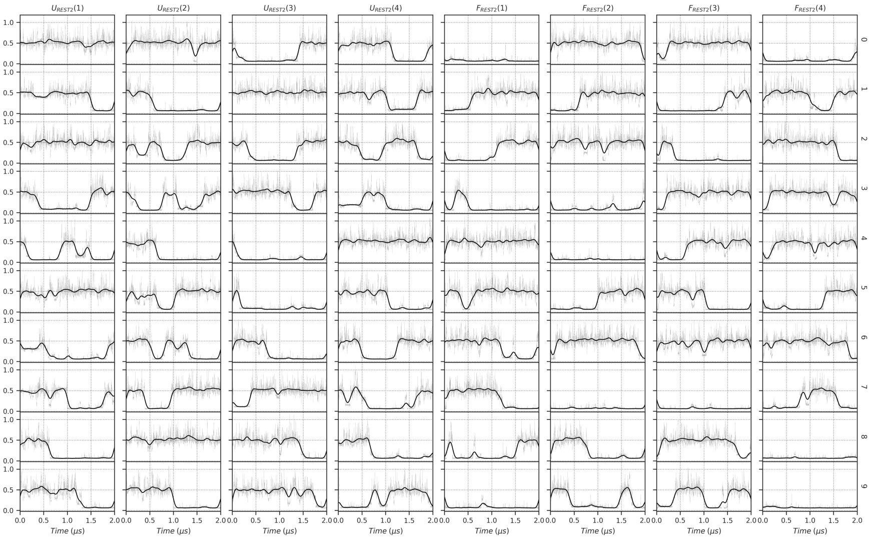

Figure S12: **RMSD as function of time for REST2 TrpCage simulations.** RMSD has been compute on backbone atoms, without the first and last residues, to reference structure 5awl.[1] Raw RMSD values are displayed as gray line, and a smooth gaussian filter with $\sigma = 20$ is applied, and displayed as black line.

#### Effect of Proline *cis-trans* transition on sampling

While $\lambda$ or temperature generally oscillated frequently between minimum and maximum values, some simulations shown a bias towards high temperature values. For example, in the CLN025 $F_{ST}(3)$ simulation, the RMSD showed an unfolded conformation for the last 3 $\mu s$ of the simulation. Similar behaviour was observed for CLN025 $F_{6\,rungs}(1)$, $F_{6\,rungs}(2)$ and $F_{350K}(3)$ simulations (Fig. S5 and S6). Visual inspection revealed that the protein was trapped in an unfolded conformation due to the proline 4 in the *cis* conformation. As shown in Fig. S13, the proline 4 was primarily in a *trans* conformation, but in rare cases, a *cis-trans* transition occurred, trapping the protein in an unfolded conformation for up to 4 $\mu s$. Fig. 3 shows that during the simulation $F_{6\,rungs}(1)$, the proline 4 in a *cis* conformation, prevented CLN025 to reach the folded state. For Trp-Cage in all but one ST and SST1 simulations, as shown in Fig. S15, at least one of the proline $\omega$ angles switched to *cis* conformation, with very heterogeneous transition time, forming few hundreds of $ns$ to more than 30.0 $\mu s$ as in Trp-Cage $F_{ST}(4)$.

The exclusion of $\omega$ dihedral angles from the solute-scaled intramolecular energy term in replicas 3 and 4 of simulations $F$ and $U$ of Trp-Cage, and in all replicas of Trp-Cage $F_{350K}$ prevents, as expected, the proline residues from switching to the *cis* conformation (Fig. S15).

he fact that a proline residue can be trapped in the *cis* conformation for more than 30.0 $\mu s$ during an ST simulation heating up the system to 500 $K$ is quite a surprise. This *cis-trans* transition is a slow process that strongly affects the sampling of the protein conformational space. As it is rare, we excluded simulation frames with proline residues in *cis* conformation from subsequent analyses comparing SST2 and to other sampling algorithm efficiency.

Given the extreme slowness of the *cis-trans* transition of proline residues, we clearly observe the advantage of excluding proline $\omega$ dihedral angles from the solute-scaled intramolecular energy term to efficiently sample the fold-unfold transition of proteins.

owever, one might be interested in sampling the conformation of proline residues for specific applications. To investigate proline *cis-trans* transition, we also performed SST2 simulations of Trp-Cage using higher range of temperature, from 280 $K$ to 700 $K$ ($F_{280-700K}$). As shown in Fig. S15 the proline residues were observed to exchange conformation with an order of magnitude more frequently, the average transition time was $129 \pm 46$ $ns$ , as for ST, SST1, REST2 and SST2 simulations, the average transition time were all greater than 400 $ns$, reaching $5.1 \pm 4.0$ and $3.1 \pm 1.6$ $\mu s$ for ST and SST2 simulation, respectively (Fig. S17).

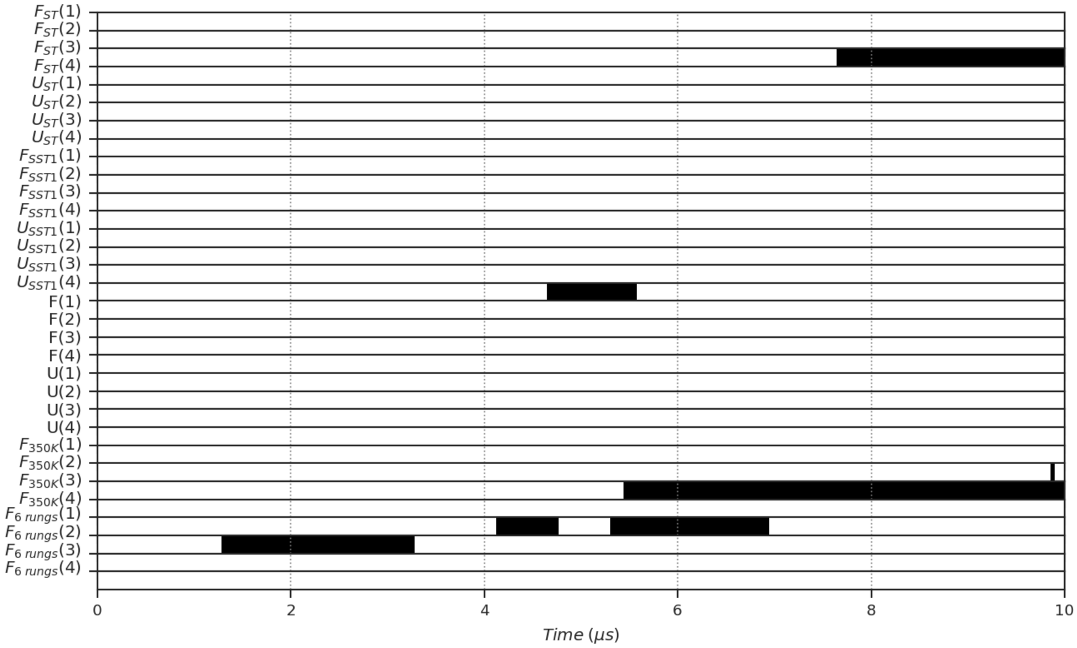


Figure S13: **CLN025 Proline 4 *cis-trans* conformation.** Proline 4 state (*cis-trans*) as function of simulation time. For each simulation, the Proline 4 residue state is displayed as a code bar, with white and black color representing Proline 4 in *trans* state and *cis* state, respectively. $\omega$ angle is defined as the dihedral angle between the $C_{\alpha(i-1)}$-$C_{(i-1)}$-$N_{(i)}$-$C_{\alpha(i)}$ atoms, where $(i)$ is the Proline residue index. $\omega$ angle is defined in *cis* state when $\omega$ angle is between -90 and 90°, and in *trans* state otherwise.

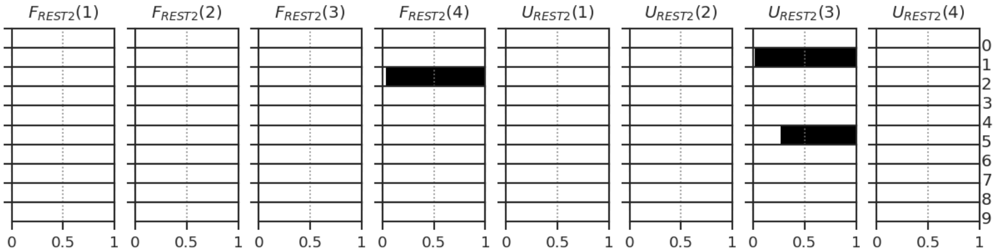


Figure S14: **CLN025 Proline 4 *cis-trans* conformation for REST2 simulations.** Proline 4 state (*cis-trans*) as function of simulation time. For each simulation, the Proline 4 residue state is displayed as a code bar, with white and black color representing Proline 4 in *trans* state and *cis* state, respectively. $\omega$ angle is defined as the dihedral angle between the $C_{\alpha(i-1)}$-$C_{(i-1)}$-$N_{(i)}$-$C_{\alpha(i)}$ atoms, where $(i)$ is the Proline residue index. $\omega$ angle is defined in *cis* state when $\omega$ angle is between -90 and 90°, and in *trans* state otherwise.

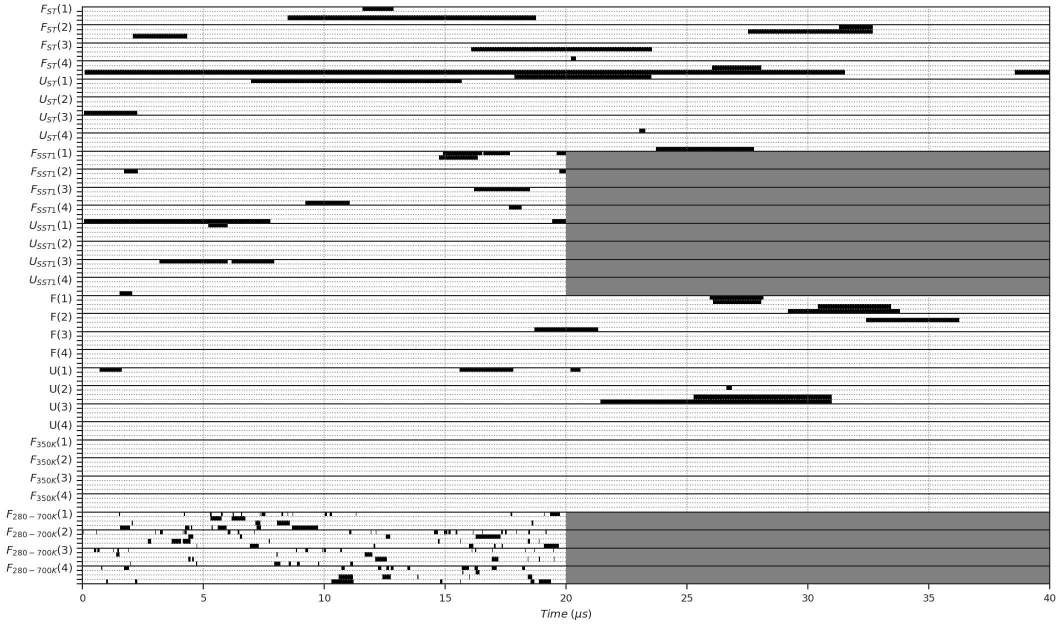


Figure S15: **Trp-Cage Proline residues *cis-trans* conformation.** Proline residues state (*cis-trans*) as function of simulation time. For each simulation, the Proline 12, 17, 18, 19 residues state are displayed as a code bar, with white and black color representing Proline *trans* state and *cis* state, respectively. $\omega$ angle is defined as the dihedral angle between the $C_{\alpha(i-1)}$-$C_{(i-1)}$-$N_{(i)}$-$C_{\alpha(i)}$ atoms, where $(i)$ is the Proline residue index. $\omega$ angle is defined in *cis* state when $\omega$ angle is between -90 and 90°, and in *trans* state otherwise.

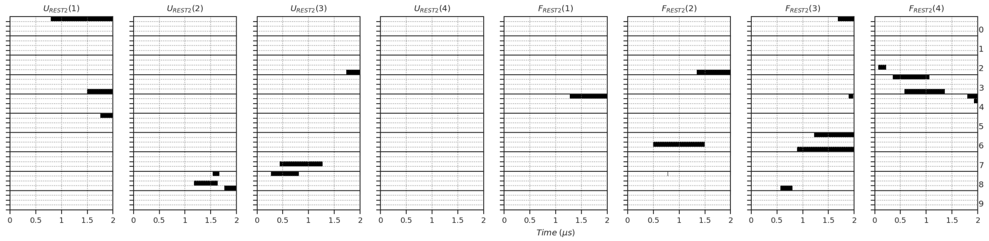


Figure S16: **Trp-Cage Proline residues *cis-trans* conformation for REST2 simulations.** Proline residues state (*cis-trans*) as function of simulation time. For each simulation, the Proline 12, 17, 18, 19 residues state are displayed as a code bar, with white and black color representing Proline *trans* state and *cis* state, respectively. $\omega$ angle is defined as the dihedral angle between the $C_{\alpha(i-1)}$-$C_{(i-1)}$-$N_{(i)}$-$C_{\alpha(i)}$ atoms, where $(i)$ is the Proline residue index. $\omega$ angle is defined in *cis* state when $\omega$ angle is between -90 and 90°, and in *trans* state otherwise.

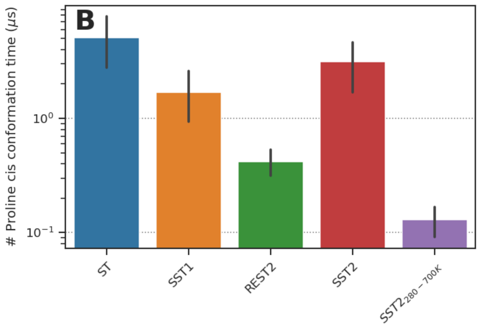


Figure S17: **Trp-Cage Proline residues *cis* persitence time.** REST2 persistence time of Proline residues in *cis* state is probably underestimated due to the fact that REST2 simulations are shorter than SST2 simulations. In REST2 simulations, out of 22 *cis* conformation transitions, 13 were still in *cis* conformation at the end of the simulation. $\omega$ angle is defined as the dihedral angle between the $C_{\alpha(i-1)}$-$C_{(i-1)}$-$N_{(i)}$-$C_{\alpha(i)}$ atoms, where $(i)$ is the Proline residue index. $\omega$ angle is defined in *cis* state when $\omega$ angle is between -90 and 90°, and in *trans* state otherwise.

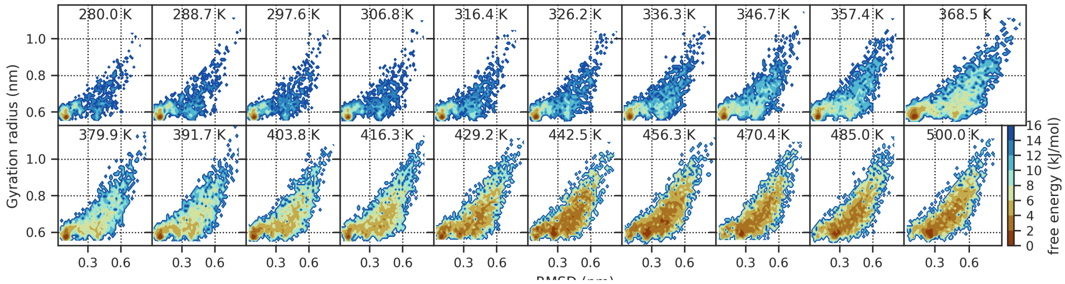


Figure S18: **Explored structural space as function of temperature for ST simulation of CLN025 $F(1)$.** Free Energy landscape as function of RMSD and radius of gyration for CLN025 $F(1)$ SST2 simulation. The color code corresponds to the free energy value in $kJ/mol$.

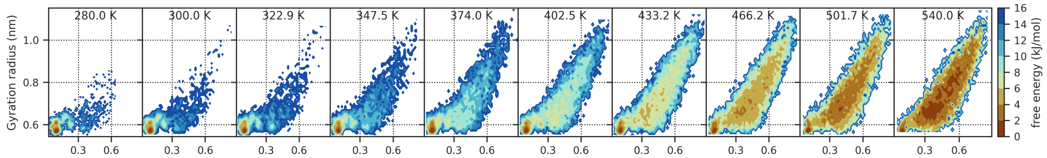


Figure S19: **Explored structural space as function of temperature for SST2 simulation of CLN025 $F(1)$.** Free Energy landscape as function of RMSD and radius of gyration for CLN025 $F(1)$ SST2 simulation. The color code corresponds to the free energy value in $kJ/mol$.

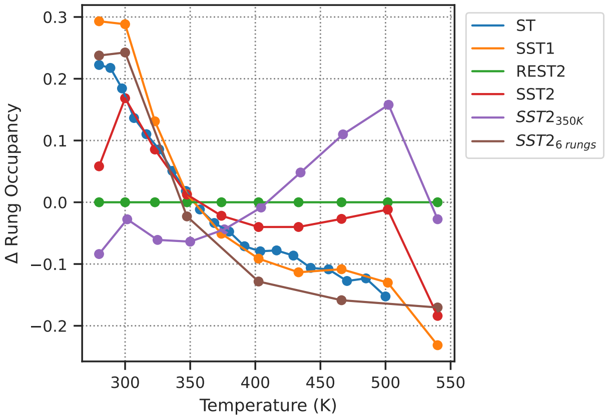


Figure S20: **CLN025 Temperature rung occupancy.** Ratio of effective occupancy to theoretical occupancy. For SST2 and REST2 simulations, the rung temperatures correspond to the solute-solute corresponding temperatures.

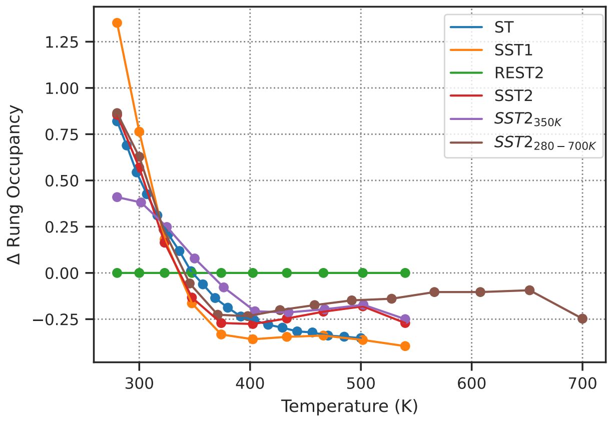


Figure S21: **Trp-Cage Temperature rung occupancy.** Ratio of effective occupancy to theoretical occupancy. For SST2 and REST2 simulations, the rung temperatures correspond to the solute-solute corresponding temperatures.

### 2 PNGase

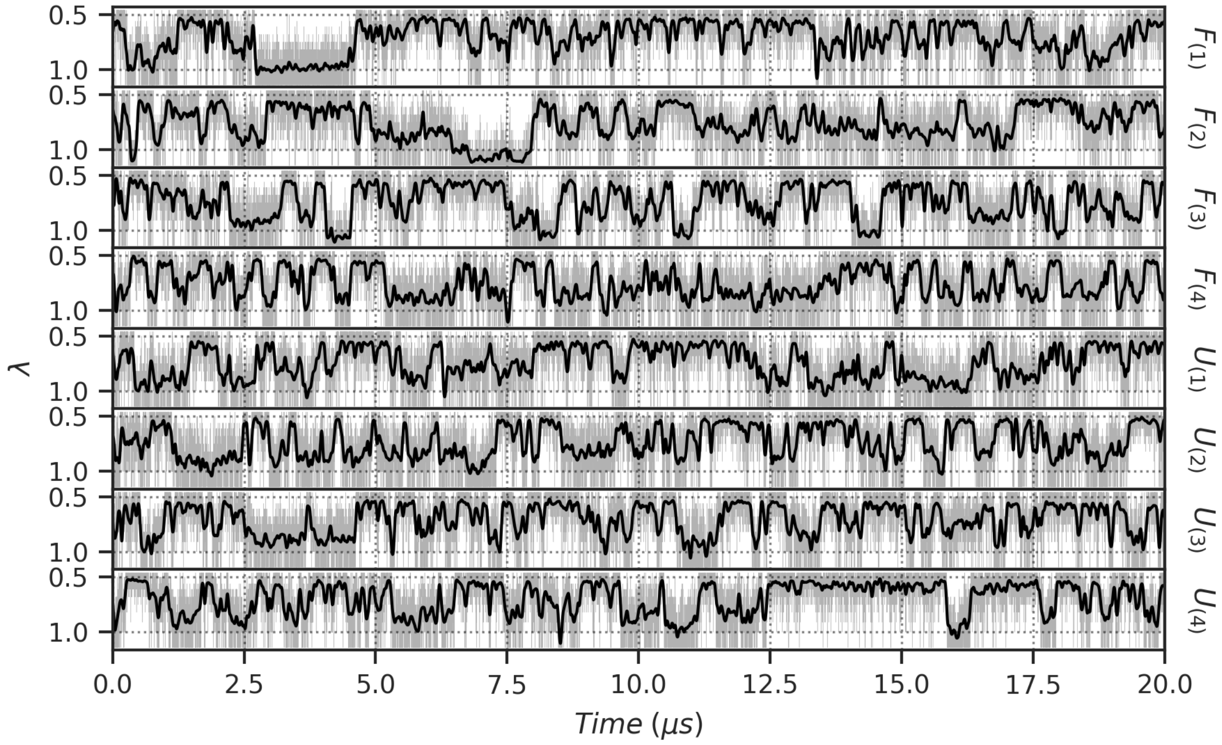


Figure S22: **$\lambda$ as function of time for PNGase simulations.** Raw $\lambda$ values are displayed as gray line, and a smooth gaussian filter with $\sigma = 20$ is applied, and displayed as black line.

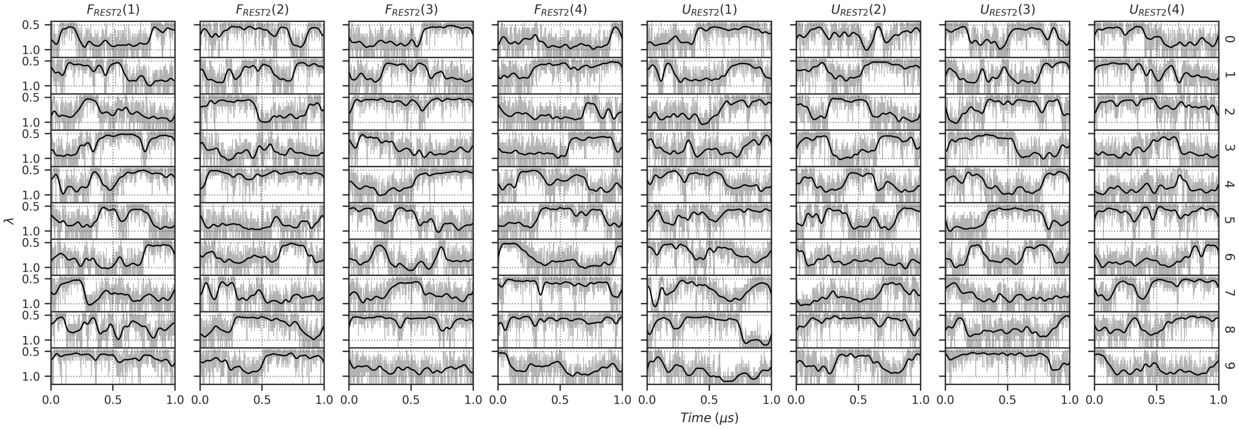


Figure S23: **λ as function of time for REST2 PNGase simulations.** Raw λ values are displayed as gray line, and a smooth gaussian filter with $\sigma = 20$ is applied, and displayed as black line.

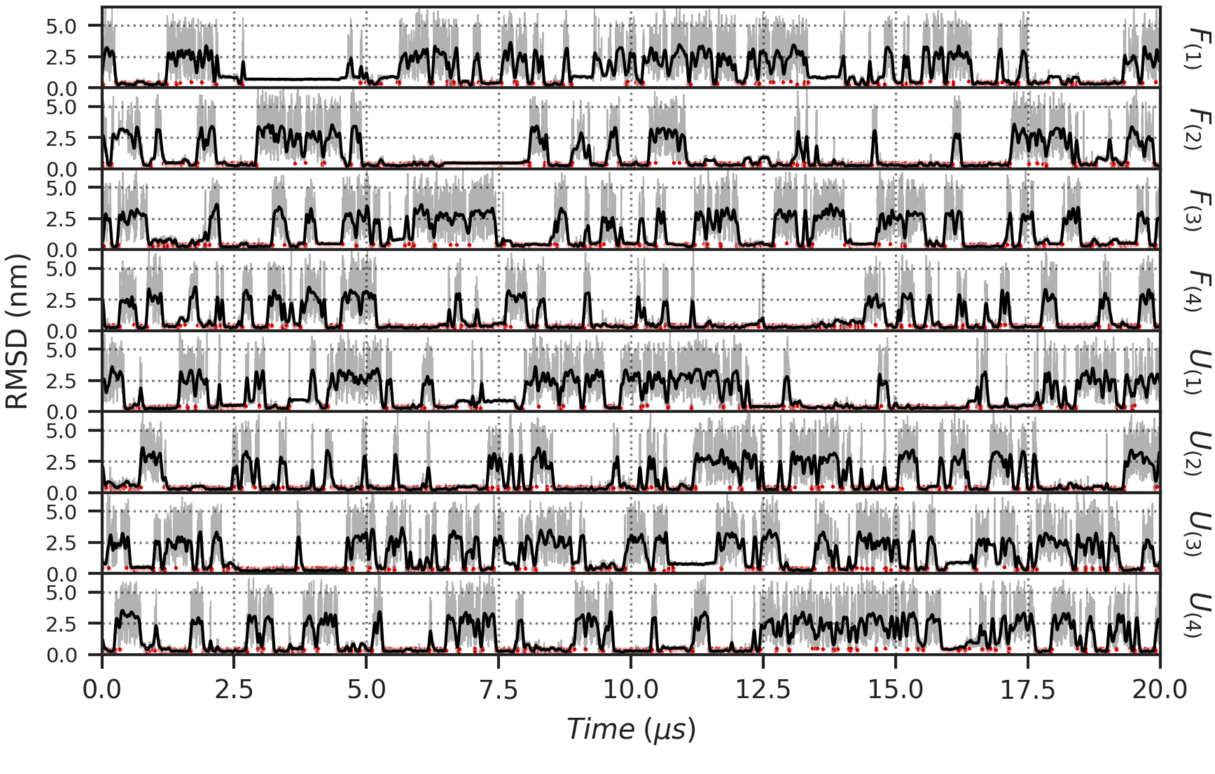


Figure S24: **RMSD as function of time for p97 peptide.** RMSD has been compute on backbone atoms of p97 peptide, after structure alignement of PNGase backbone atoms, to reference structure 2hpl.[3] Raw RMSD values are displayed as gray line, and a smooth gaussian filter with $\sigma = 20$ is applied, and displayed as black line. Frames in which the peptide is considered as bounded (RMSD < 0.5 $nm$) are indicated as red dots.

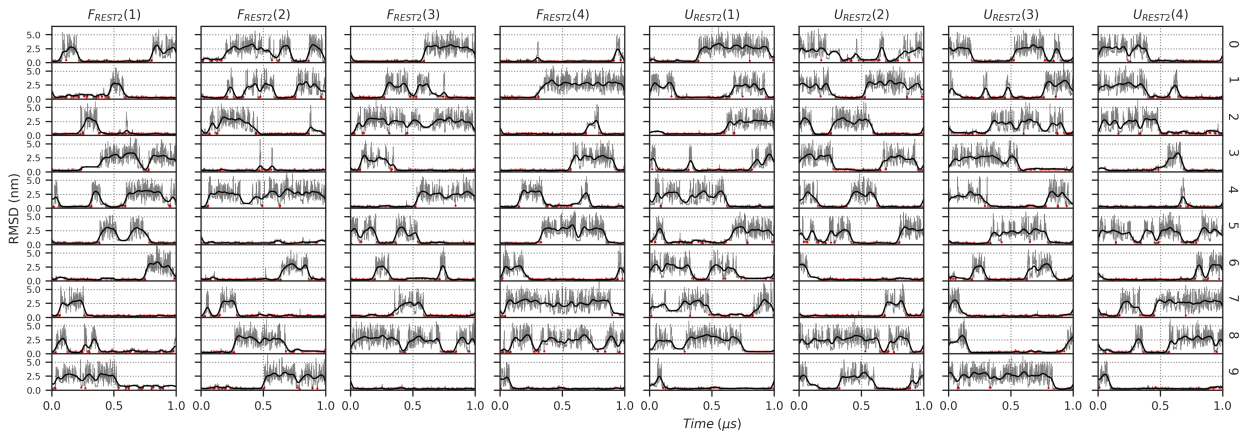


Figure S25: **RMSD as function of time for p97 peptide during REST2 simulations.** RMSD has been compute on backbone atoms of p97 peptide, after structure alignement of PNGase backbone atoms, to reference structure 2hpl.[3] Raw RMSD values are displayed as gray line, and a smooth gaussian filter with $\sigma = 20$ is applied, and displayed as black line. Frames in which the peptide is considered as bounded (RMSD $< 0.5$ $nm$) are indicated as red dots.

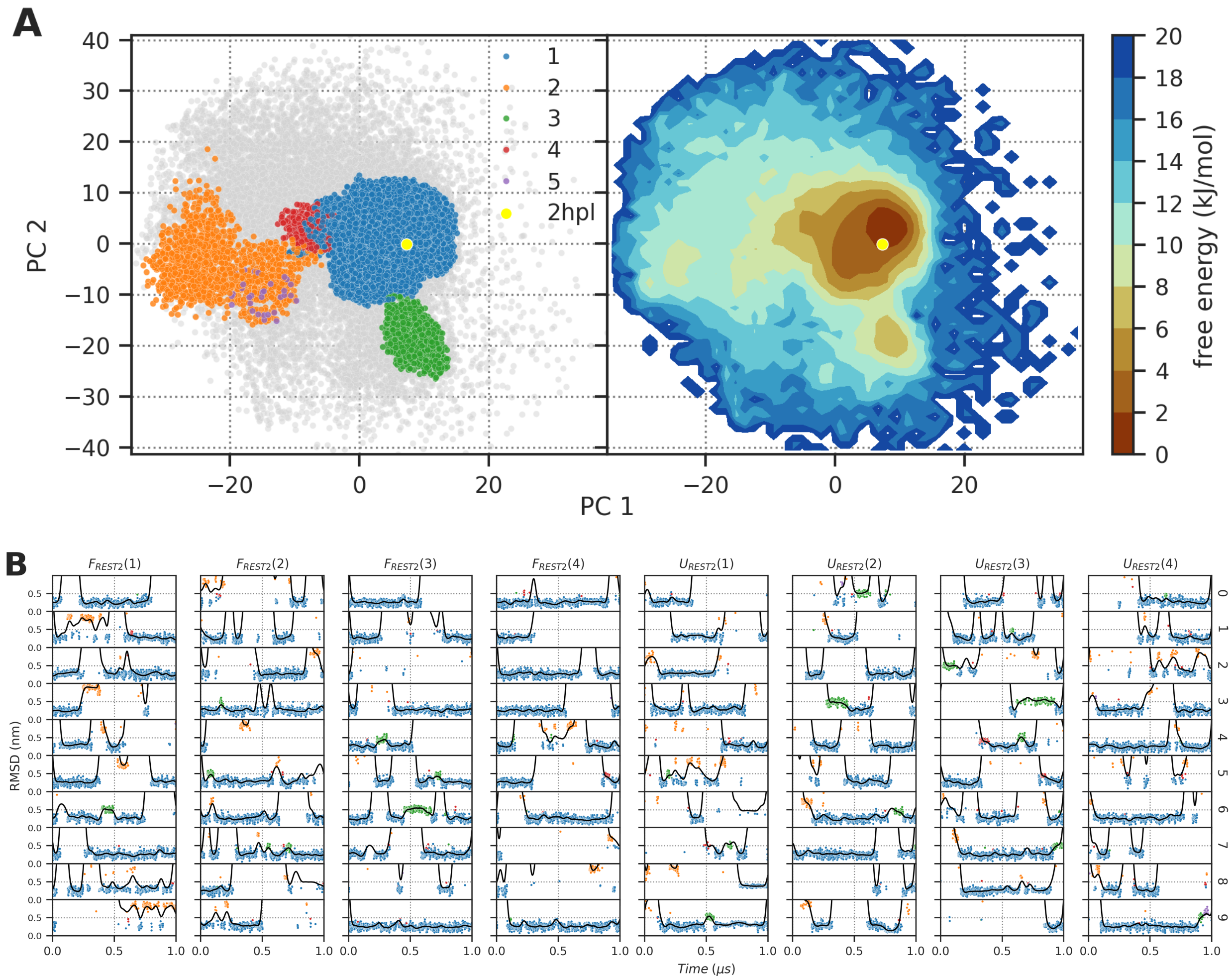


Figure S26: **Peptide conformation in the binding site during REST2 simulations.** **(A)** Projection of the two first principal component of p97 backbone atoms computed after trajectory alignement on PNGase backbone atoms. Only simulation frames where the peptide is in the binding site are displayed (peptide RMSD lower than 1.0 $nm$). Left panel displays the projection of clustered simulation frames colored by clusters, non clustered frames are displayed as gray points. Right panel displays the free energy landscape computed from the projection of the binding site simulation frames. The reference structure 2hpl[3] conformation is displayed as a yellow dot. **(B)** RMSD as function of time for p97 peptide. RMSD has been compute on backbone atoms of p97 peptide, after structure alignement of PNGase backbone atoms, to reference structure 2hpl. Raw RMSD values are displayed as dots, colored as function of clusters, blue, orange, green, red and purple for clusters 1, 2, 3, 4 and 5, respectively. A smooth gaussian filter with $\sigma = 20$ applied on RMSD values, is displayed as black line.

### 3 Simulations convergence

To assess the convergence of weight estimation for SST1, SST2 and ST simulations, it is first necessary to assess the distribution of the explored $\lambda$ and temperature space. As illustrated in Fig. S20 and S21, the rung occupancy of the SST2 simulations is comparable to that of the ST simulations. In the case of CLN025, the distribution of explored temperature space is mostly biased towards low temperature as evidenced by the $ST$, $SST1$, $SST2$ and $F_{6\ rungs}$ simulations. The latter group of simulation exhibited over-exploration up to 30% at 280 $K$. The only exception is the $SST2_{350\ K}$ simulation, which exhibited moderate over-exploration at high temperature, up to 15% at 500 $K$. With regard to Trp-Cage, the distribution of explored temperature space is relatively homogeneous, with for all simulations a surexploration of low temperature rungs up to 130% at 280 $K$, and about down to -40% at 500 $K$.

Fig. S27 illustrates the convergence of weight estimation RMSD for SST1, SST2 and ST simulations. As the ST simulations do not have the same number of water molecules and atoms, a direct comarison of the $E_p$ values between replicas is not possible. Therefore, the RMSD of the $w_0, w_1, \ldots, w_{i-1}$ weight values was computed separately for each replica and averaged. For SST1 and SST2, the final weight values were computed as the average of all SST1 and SST2 replicas and simulations, and therefore did not converge to zero.

As illustrated in Fig. S27A and S27B, the weight estimation convergence is comparable between SST1, SST2 and ST simulations. The weight estimation convergence for CLN025 is reached after approximately 2 $\mu s$ of simulation, while for Trp-Cage, it is reached after approximately 5–10 $\mu s$ of simulation. With regard to the PNGase/p97 complex, the weight estimation convergence is attained after approximately 2–3 $\mu s$ of simulation (Fig. S27C).

Furthermore, the convergence of the folding fraction by temperature for CLN025 and Trp-Cage was computed as a function of simulation time, as was the convergence of the fraction of peptide bound by temperature for the PNGase/p97 complex. As illustrated in Fig. S28, the convergence is faster for the folded or bound fraction RMSD than for the weight estimate RMSD. The convergence of the folded fraction for CLN025 and Trp-Cage is reached after approximately 0.5 $\mu s$ and 5 $\mu s$ respectively. The convergence of the bound fraction for the PNGase/p97 complex is reached after approximately 1 $\mu s$ of simulation. It should be noted that the elevated error bar observed for Trp-Cage $ST$ simulations can be attributed to the fact that the protein was confined to an unfolded state for a significant portion of the simulation, exceding 80 % of the total simulation time in the $F_{ST}(4)$ simulation. As observed in the $F(1)$ simulation of the PNGase/p97 complex, the peptide remained in unbound conformations for extended periods of time, which notably impacted the fraction of peptide bound as a function of temperature for $SST2$ simulations.

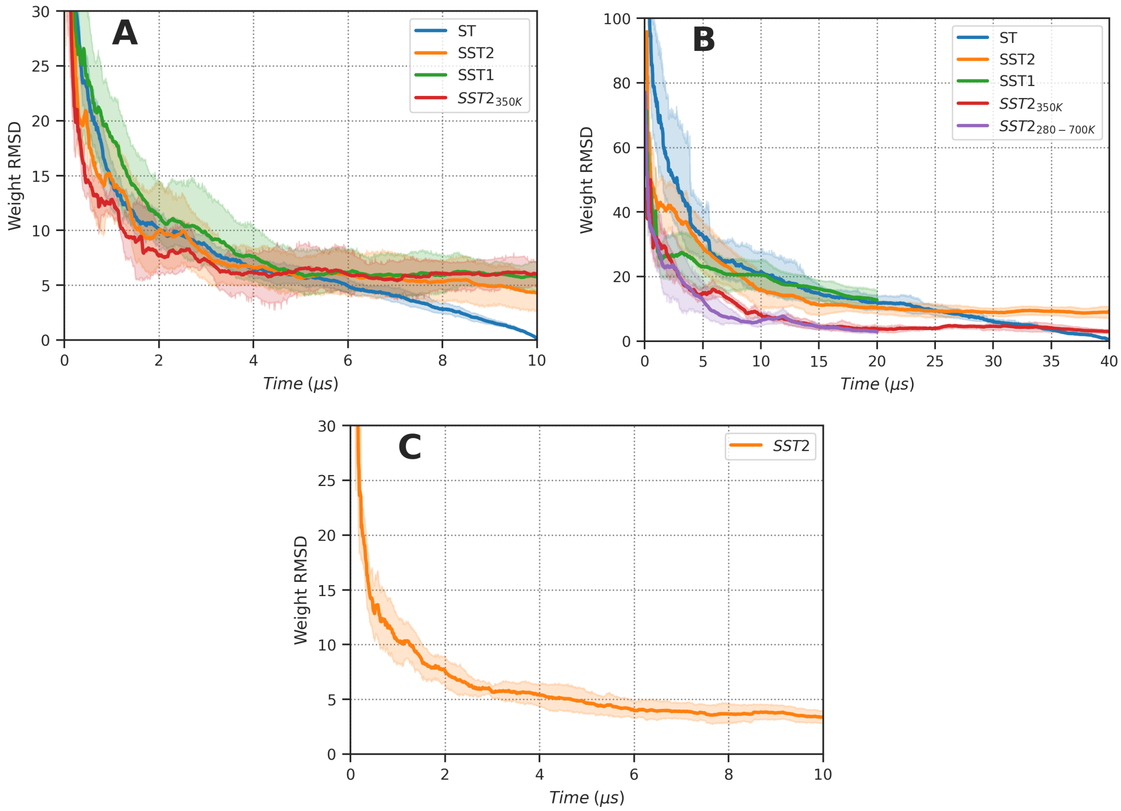


Figure S27: **Convergence of weight estimation.** RMSD of $w_0, w_1, \ldots, w_{i-1}$ weights values compare to final ones as function of simulation time for CLN025 **(A)**, Trp-Cage **(B)** and PNGase/p97 complex **(C)**. Note that as ST simulations do not have the same number of water molecules and of atoms, the $E_p$ value cannot be compared between replicas. Consequently RMSD of $w_0, w_1, \ldots, w_{i-1}$ weights values have been computed separately for each replicas and averaged, this explain why all ST RMSD converge to zero. For SST1 and SST2, the final weight values were computed as the average of all SST1 and SST2 replicas and simulation, and thus do not converge to zero. For all panels, transparent layer around the plain line displays the 95 % confidence interval computed over the 4 replicas.

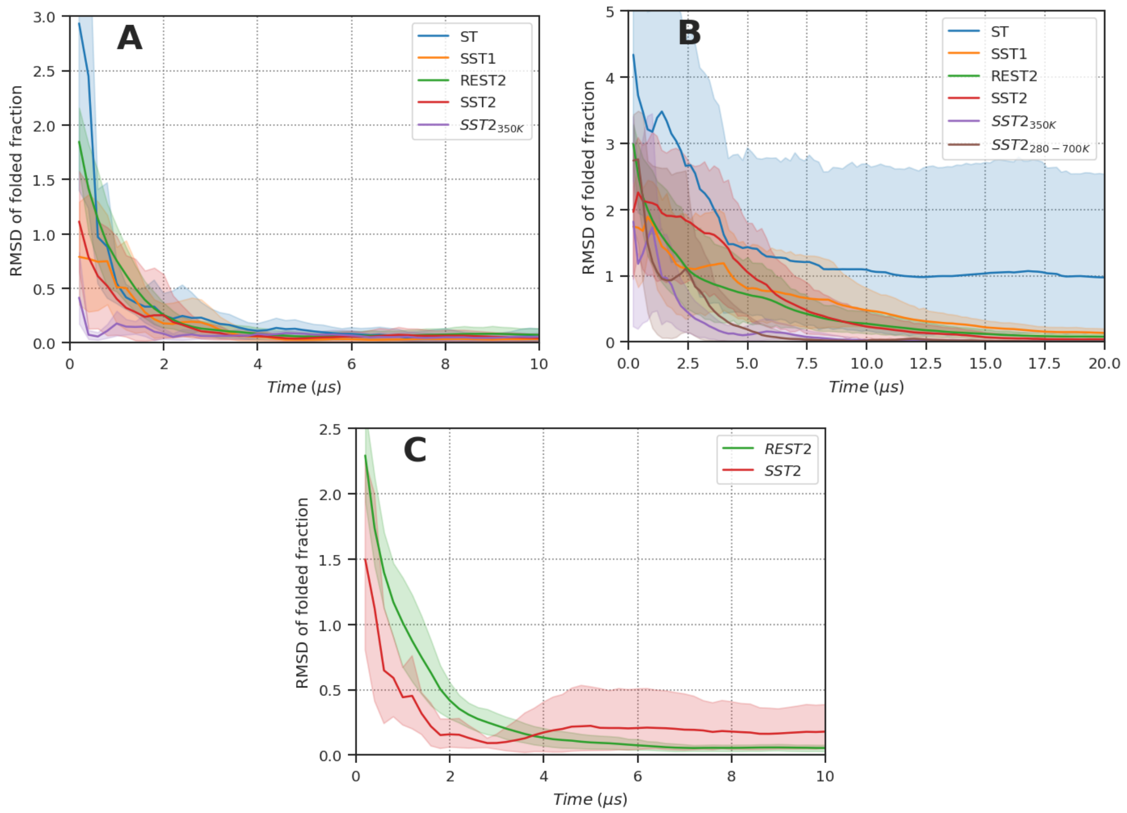


Figure S28: **ST, SST1, SST2 and REST2 stability curve convergence.** RMSD of folding fraction by temperature for CLN025 **(A)** and Trp-Cage **(B)**. **(C)** RMSD of bound peptide fraction by temperature for PNGase-p97 complex. Every 200 $ns$, the fraction of folded or bound peptide is computed and the RMSD to the final stability curve is computed. For all panels, transparent layer around the plain line displays the 95 % confidence interval computed over the 4 replicas.

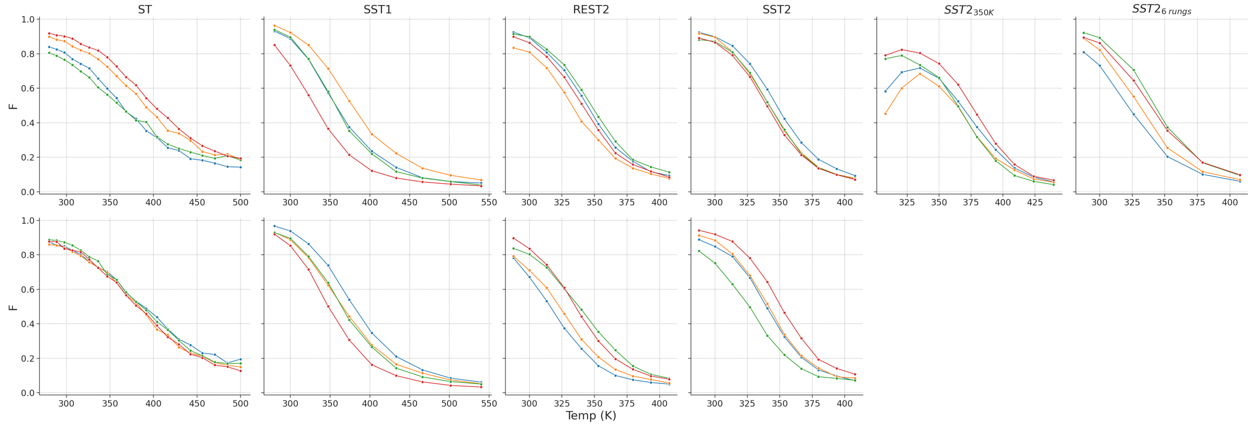


Figure S29: **Folded fraction details of CLN025.** Fraction of folded conformation as function of temperature for CLN025 simulations. Simulation frames with Proline $\omega$ angle in *cis* conformation have been excluded to compute fraction of folded conformation. CLN025 was considered folded when its RMSD was below 0.2 *nm*. Upper panel displays simulations starting from the Folded state, and lower panel displays simulations starting from the Unfolded state.

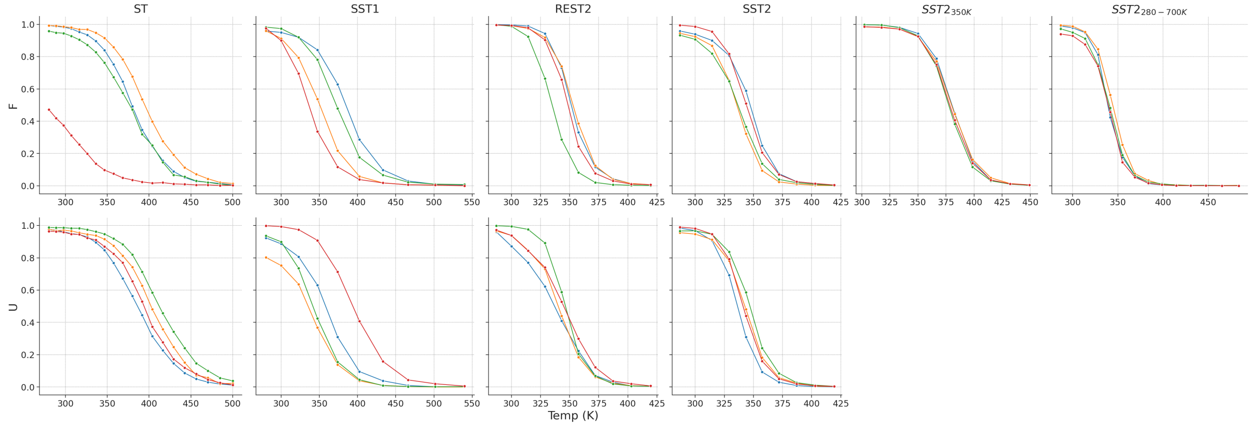


Figure S30: **Folded fraction details of TrpCage.** Fraction of folded conformation as function of temperature for TrpCage simulations. TrpCage was considered folded when its RMSD was below 0.2 *nm*. Upper panel displays simulations starting from the Folded state, and lower panel displays simulations starting from the Unfolded state.

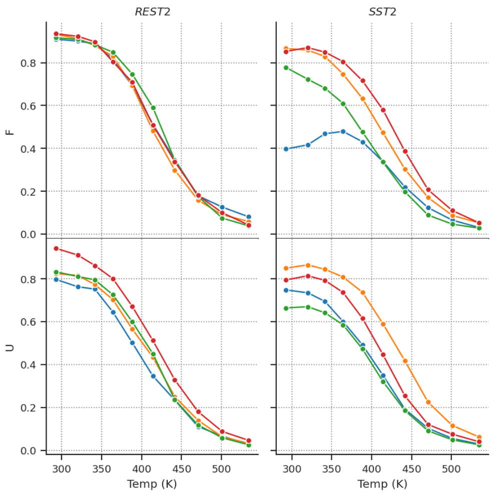


Figure S31: **Bound fraction details of PNGase.** Fraction of bounded conformation as function of temperature for PNGase/p97 simulations. p97 was considered as bound when their backbone RMSD to 2hpl was below 0.5 $nm$. Upper panel displays simulations starting from the bounded state, and lower panel displays simulations starting from the unbounded state.

### 4 Effect of solvent box size on effective temperature estimation

To directly assess whether the number of water or size of solvent box affect the effective melting temperature estimation, we performed SST2 simulations using a larger solvent box. This increases the number of water molecules and thus alters the solute–solvent interaction energy. As shown in Figure S32A, the resulting melting temperatures remain effectively unchanged. Interestingly, we observe that a decrease in solute–solvent interaction energy (Fig. S32B) appears to be compensated in part by an increase in solute–solute interaction energy (Fig. S32C), leading to stable overall behavior. This suggests that the size of solvent box in the SST2/REST2 framework is not affecting the $T_m$ estimation.

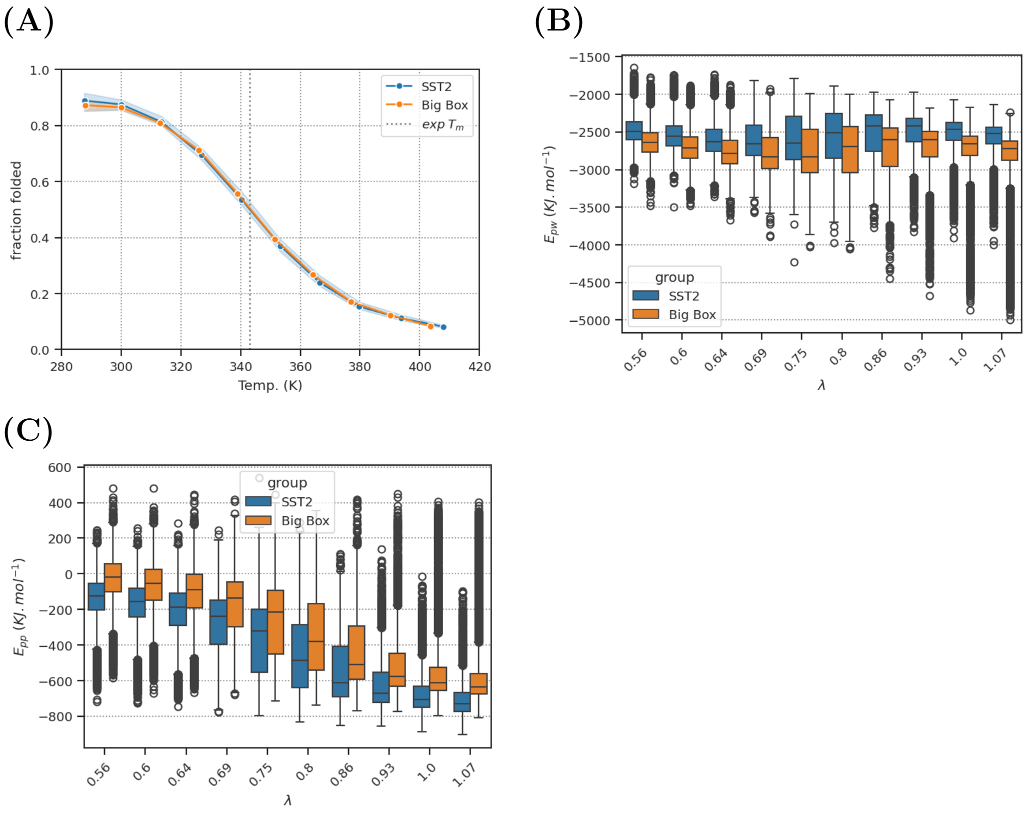


Figure S32: **Effect of simulation box size on folded fraction as function of temperature for TrpCage.** **(A)** Folded fraction as function of temperature for TrpCage SST2 simulations with classical simulation box (1.5 $nm$ padding) and larger simulation box (3.0 $nm$ padding). The folded fraction is computed as the fraction of frames with RMSD below 0.2 $nm$ to the reference structure 1l2y.[2] **(B)** Solute-solvent interaction energy as function of lambda for TrpCage SST2 simulations with classical simulation box and larger one. **(C)** Solute-solute interaction energy as function of lambda for TrpCage SST2 simulations with classical simulation box and larger one.

### 5 SST2 computing performance

In order to accurately compute the long-range electrostatic contribution of $E_{pp}$ and $E_{ww}$, it is necessary to create two additional systems whose coordinates are assigned as the simulated system each time a $\lambda$ exchange is attempted. This process is time-consuming and results in a moderate slowdown when using the SST2 simulation, with a reduction of efficiency of approximately 20 % compared to a conventional simulation. The simulation of the mouse p97/PNGase complex (PDBID:2HPL[3]) solvated in water (14,867 atoms) using an integration time step of 4 $fs$ demonstrated a performance of 697.7 $ns/day$ on an NVIDIA RTX A4000 GPU card. The separation of the energy terms (REST2 implementation) resulted in a performance of 612.7 $ns/day$, representing a slowdown of approximately 12 %, while running SST2 with a temperature change interval of 2 $ps$ yielded a performance of 571.6 ns/day, indicating a slodown of approximately 18 % compared to conventional MD simulation.

### 6 Temperature swap interval

In both SST2 and ST simulations, a systematic evaluation of the requisite time interval for temperature swap was conducted. Short simulations for SST2 and ST were carried out with a time interval of 10 $ps$, with energy terms saved every 0.1 $ps$. The SST2 and ST simulations were 1.0 $\mu s$ and 500 $ns$ long, respectively (see Table S1).

To evaluate the time interval, three temperature ladders were uemployed, comprising 280.0, 288.4, and 297.0 $K$ for ST and $\lambda$ values of 1.0, 0.935, and 0.875 for SST2, with a reference temperature of 280 $K$. The aforementioned $\lambda$ values correspond to solute-solute temperatures of 280.0, 299.3, and 320.0 $K$.

During SST2 simulations, the distribution of the energy terms $E_{pp} + 0.5\sqrt{\beta_{ref}/\beta_m}E_{pw}$ was measured at 0.5, 1.0, 2.0, and 4.0 $ps$ after temperature swap (Fig. S33A). The distribution rapidly converged and achieved canonical behaviour within a timsescale of 0.5 to 1 $ps$. In contrast, convergence of potential energy was observed to occur at a slower rate in ST simulations. Fig. S33B and S33C illustrate that, at 2.0 $ps$, the energy difference had not yet converged; only at 4.08.0 $ps$ could the potential energy distribution be considered canonical. This is in contrast with the findings presented in,[4] where faster convergence after a temperature swap was observed. This discrepency may be attributed to differences in the thermostat utilised in MD simulations. To ensure consistent exchange times between SST2 and ST, the friction coefficient of the Langevin integrator was updated from 1 to 10 $ps^{-1}$. As illustrated in Fig. S33C and S33D, the potential energy exhibited a markedly accelerated convergence when a friction coefficient of 10 $ps^{-1}$ was employed, attaining canonical status at 2$ps$. SST2 simulations were also conducted with a friction coefficient of 10 $ps^{-1}$ and no discernible differences were observed. This led to the conclusion that the exchange time should be set to 2 $ps$ for subsequent simulations (Table 1).

Table S1: List of short simulation of Chignolin CLN025. PDB reference ID structure for CLN025 is 5awl.[1]

| name | protein | starting form | length ($\mu s$) | exchange time ($ps$) | friction ($ps^{-1}$) | Sampling Method | min. Temp. | max. Temp. | ref. Temp. | #Rung | #Replicas |
|---|---|---|---|---|---|---|---|---|---|---|---|
| $F\_ST_{friction\ 1}$ | CLN025 | 5awl | 0.5 | 10.0 | 1.0 | ST | 280 | 297 | | 3 | 1 |
| $F\_ST_{friction\ 10}$ | CLN025 | 5awl | 0.5 | 10.0 | 10.0 | ST | 280 | 297 | | 3 | 1 |
| $F_{friction\ 1}$ | CLN025 | 5awl | 1.0 | 10.0 | 1.0 | SST2 | 280 | 320 | 280 | 3 | 1 |
| $F_{friction\ 10}$ | CLN025 | 5awl | 1.0 | 10.0 | 10.0 | SST2 | 280 | 320 | 280 | 3 | 1 |

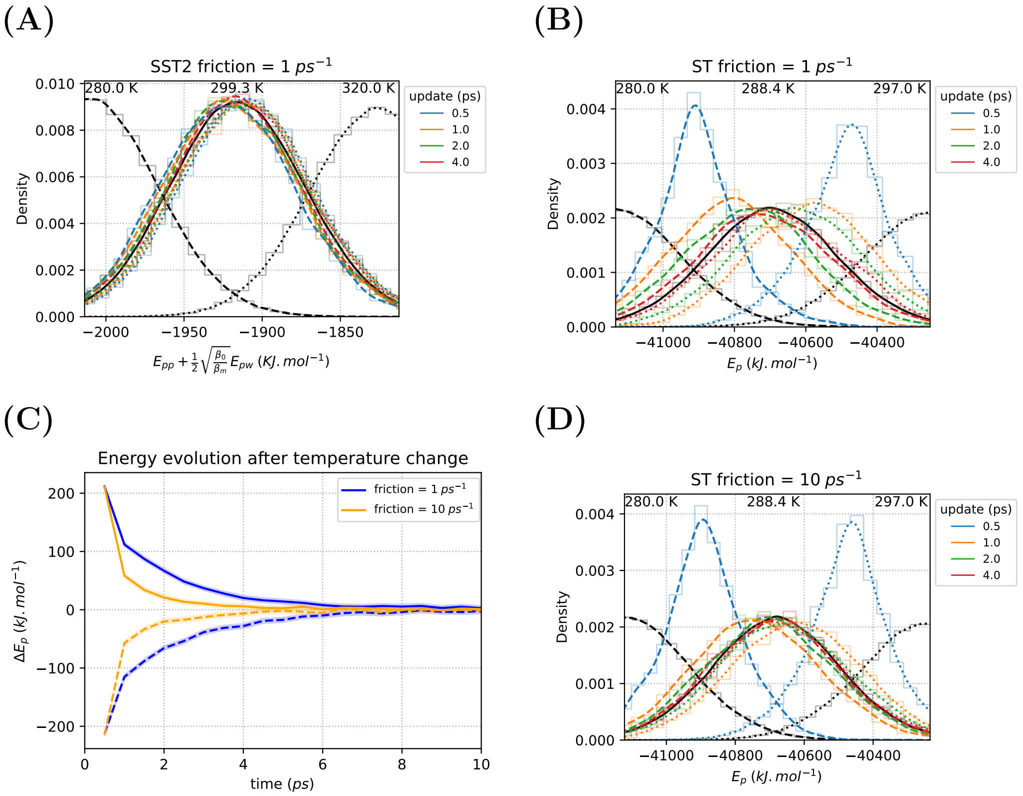


Figure S33: **Potential Energy relaxation after temperature update.** **(A)** Distribution of $E_{pp} + 1/2\sqrt{\beta_{ref}/\beta_m}E_{pw}$ after $\lambda$ update during SST2 F$_{friction1}$ simulation at different time interval. **(B)** Distribution of potential energy $E_p$ after temperature update during ST F_ST$_{friction1}$ simulation with a friction coefficient of 1 $ps$−1. **(C)** Evolution of potential energy after temperature update during ST simulations, F_ST$_{friction1}$ and F_ST$_{friction10}$, with friction coefficients of 1 and 10 $ps^{-1}$, displayed as blue and orange curve respectively. **(D)** Distribution of potential energy $E_p$ after temperature update during ST $F\_ST_{friction\ 10}$ simulation with a friction coefficient of 10 $ps$−1.
